# Bacterial genome encoded ParMs

**DOI:** 10.1101/2024.09.12.612785

**Authors:** Samson Ali, Adrian Koh, David Popp, Kotaro Tanaka, Yoshihito Kitaoku, Naoyuki Miyazaki, Kenji Iwasaki, Kaoru Mitsuoka, Robert C. Robinson, Akihiro Narita

**Affiliations:** Research Institute for Interdisciplinary Science, Okayama University, Okayama 700-8530, Japan; Institute of Molecular and Cell Biology, A*STAR (Agency for Science, Technology and Research), Biopolis, Singapore 138673, Singapore; Graduate School for Integrative Sciences and Engineering, National University of Singapore, Singapore 138632, Singapore; Cellular and Structural Physiology Institute (CeSPI), Nagoya University, Furo-cho, Chikusa-ku, Nagoya 464-8601, Japan; Division of biological science, Graduate School of Science, Nagoya University, Furo-cho, Chikusa-ku, Nagoya, 464-8601, Japan; The Laboratory of Protein Synthesis and Expression, Institute for Protein Research, Osaka University, Japan; Life Science Center for Survival Dynamics, Tsukuba Advanced Research Alliance (TARA), University of Tsukuba, Tsukuba, Japan; Research Center for Ultra-High Voltage Electron Microscopy, Osaka University, 7-1 Mihogaoka, Ibaraki, Osaka, 567-0047, Japan; School of Biomolecular Science and Engineering (BSE), Vidyasirimedhi Institute of Science and Technology (VISTEC), Rayong, 21210, Thailand

**Keywords:** DNA segregation, nucleotide hydrolysis, Plasmid, ParM, ParCMR system

## Abstract

ParMs generally exist on low copy number plasmids where they contribute to plasmid segregation and stable inheritance. We carried out bioinformatics analysis, which indicated that ParM genes are not only confined to plasmids but are also occasionally found on genomes. Here we report the discovery and characterization of two chromosome encoded ParMs (cParMs) from the genomes of Desulfitobacterium hafniense (Dh-cParM1) and Clostridium botulinum (Cb-cParM). Both cParMs form filaments, exhibit nucleotide hydrolysis, and possess characteristic ParM subunit structures. Dh-cParM1 forms single and tightly coupled double filaments and is highly conserved on the chromosomes of five of six Desulfitobacterium species. Interestingly, these bacteria have not been reported to harbour plasmids. Cb-cParM possesses unique properties. Its filaments were stable after nucleotide hydrolysis and Pi release, and its ParR, (Cb-cParR) did not affect the initial stage of Cb-cParM polymerization but displayed properties of a depolymerization factor for mature filaments. These results indicate functional, polymerizing ParMs can be encoded on genomes, suggesting that ParM roles may extend to other functions beyond plasmid segregation.

## Introduction

In contrast to the random diffusion mode of segregation for high copy number plasmids, low copy number plasmids require an active plasmid maintenance system to ensure equal distribution to daughter cells during cell division (Galkin et al., 2009; Nordström and Austin, 1989). Polymerizing ParM proteins belong to the ParCMR system, one of the three known types of the segregation or partition (par) systems that aid plasmid inheritance. ParM was originally discovered from the 100 kilobase pair multidrug resistant low-copy-number *E. coli* R1 plasmid (ParM-R1) (Campbell and Mullins, 2007; Jensen and Gerdes, 1997). Its polymerization provides the driving force to push two plasmids apart by binding to an adaptor protein, ParR which also binds to *parC*, a centromere-like DNA sequence located on the plasmid (Ebersbach and Gerdes, 2005). Thus, ParM is able to partition the two plasmids to the two extremes of the cell leading to faithful inheritance (Campbell and Mullins, 2007; Choi et al., 2008; Møller-Jensen et al., 2003; Salje and Löwe, 2008). The ParCMR systems are highly divergent, and a wide range of dynamics, nucleotide dependencies and ParM filament architectures have been observed for the small number of systems studied to date (Bharat et al., 2015; Jiang et al., 2016; Koh et al., 2019; Popp et al., 2012).

ParM homologues were initially thought to be restricted to a close group of γ-proteobacteria due to high sequence variation and the limited number of ParMs characterized at the protein level. “ParM” genes from Firmicutes and cyanobacteria were postulated to represent other classes of bacterial actin-like proteins, rather than ParM orthologues (Wickstead and Gull, 2011). The recent characterization of a number of actin-like proteins or “ParMs” encoded on plasmids of Firmicutes bacteria such as the archaeal actin-like protein, pSK41 from *Staphylococcus aureus*, BtParM, from *Bacillus thuringiensis*, Alp7A and AlfA both from *Bacillus subtills*, pCBH ParM filament from *Clostridium botulinum* have all indicated properties and features of ParMs similar to those of γ-proteobacteria (Brzoska et al., 2016; Derman et al., 2009; Jiang et al., 2016; Koh et al., 2019; Polka et al., 2009; Popp et al., 2010; Szewczak-Harris and Löwe, 2018). They all possess a closed beta-barrel domain, a distinguishing feature of ParMs and could form filaments in a nucleotide dependent manner as well as the presence of ParR and *parC* near the ParM sequence (Koh et al., 2019), which led to the realization that ParMs are widespread and diverse in bacteria.

More than 65% of all sequenced bacterial genomes possess a chromosomally encoded partitioning (*par*) locus (Livny et al., 2007). This *par* system (ParABS) encodes chromosomal analogs of the active plasmid partitioning systems (ParCMR), which are involved in chromosome segregation (Ptacin et al., 2010). The ParABS system has been directly implicated in the chromosome segregation in *B. subtilis* and *Caulobacter crescentus* (Lewis and Errington, 1997; Mohl and Gober, 1997). However, ParABS systems are not present in many bacteria such as enteric bacteria, including *E. coli*, *Salmonella* sp., *Haemophilus* sp., and *Yersinia* sp (Livny et al., 2007). Thus, it is possible that undiscovered active chromosome segregation systems are yet to be discovered. During our sequence searches for novel plasmid ParCMR systems, we noticed that several of these systems are encoded on chromosomes rather than on plasmids. Characterization at the molecular level of chromosome encoded ParCMR systems (cParCMR) has yet to be explored. Here we report the presence and characteristics of novel cParMs from the genomes of *Desulfitobacterium species* and *Clostridium botulinum*.

## Results

### Putative cParCMR systems

From Blast sequence searches (Camacho et al., 2009), we identified many putative cParCMR systems encoded on chromosomes (**Fig. 1**). These were located by searching for actin-fold signature sequences against fully sequence genomes and inspecting up- and down-stream for potential ParR-like protein sequences and *parC*-like DNA palindromic repeat motifs. We found systems which appear to lack cParR, such as the *Natranaerobius thermophilus* strain JW/NM-WN-LF *Nt*-cParCMR-2 cassette, and others contain more than one putative cParR or c*parC*, such as the *Bacillus tropicus* strain FDAARGOS_920 *Bt*-cParCMR-2 cassette. Some chromosomes encode more than one potential cParCMR system, as seen for *Desulfitobacterium hafniense* strain Y51 *Dh*-cParCMR-1,2,3 cassettes, *Natranaerobius thermophilus* strain JW/NM-WN-LF 1 *Nt*-cParCMR-1,2 cassettes, and *Bacillus tropicus* strain FDAARGOS_920 *Bt*-cParCMR-1,2 cassettes. *Nt*-cParM was not observed to be conserved on chromosomes from other *Natranaerobius* species, however *Bt*-cParM and *Mt*-cParM (from *Moorella thermoacetica* strain 39073-HH) are conserved on several *Bacillus* and *Moorella* species chromosomes, respectively (**Table S1-S3**). Some bacterial strains encoding cParMs (**Fig. 1**), such as *Desulfitobacterium* and *Moorella* species, do not habour plasmids (Bengelsdorf et al., 2015; Nonaka et al., 2006) whereas plasmids are known for other species, such as *Natranaerobius thermophilus* strain JW/NM-WN-LF. *Cb*-cParCMR was located on the whole genome shotgun sequences or complete chromosomes of *Clostridium botulinum* with genome sizes ranging from 3.9 to 4.3 mb. We confirmed the ParM fold of the cParMs by calculating the AlphaFold2 (Senior et al., 2020) predicted structures and inspecting them for the closed beta-barrel domain architecture (**Fig. S1**). We proceeded to characterize two cParCMR systems, *Dh*-cParCMR-1 and *Cb*-cParCMR, to ascertain whether they have similar properties to plasmid encoded systems.

**Figure 1:**
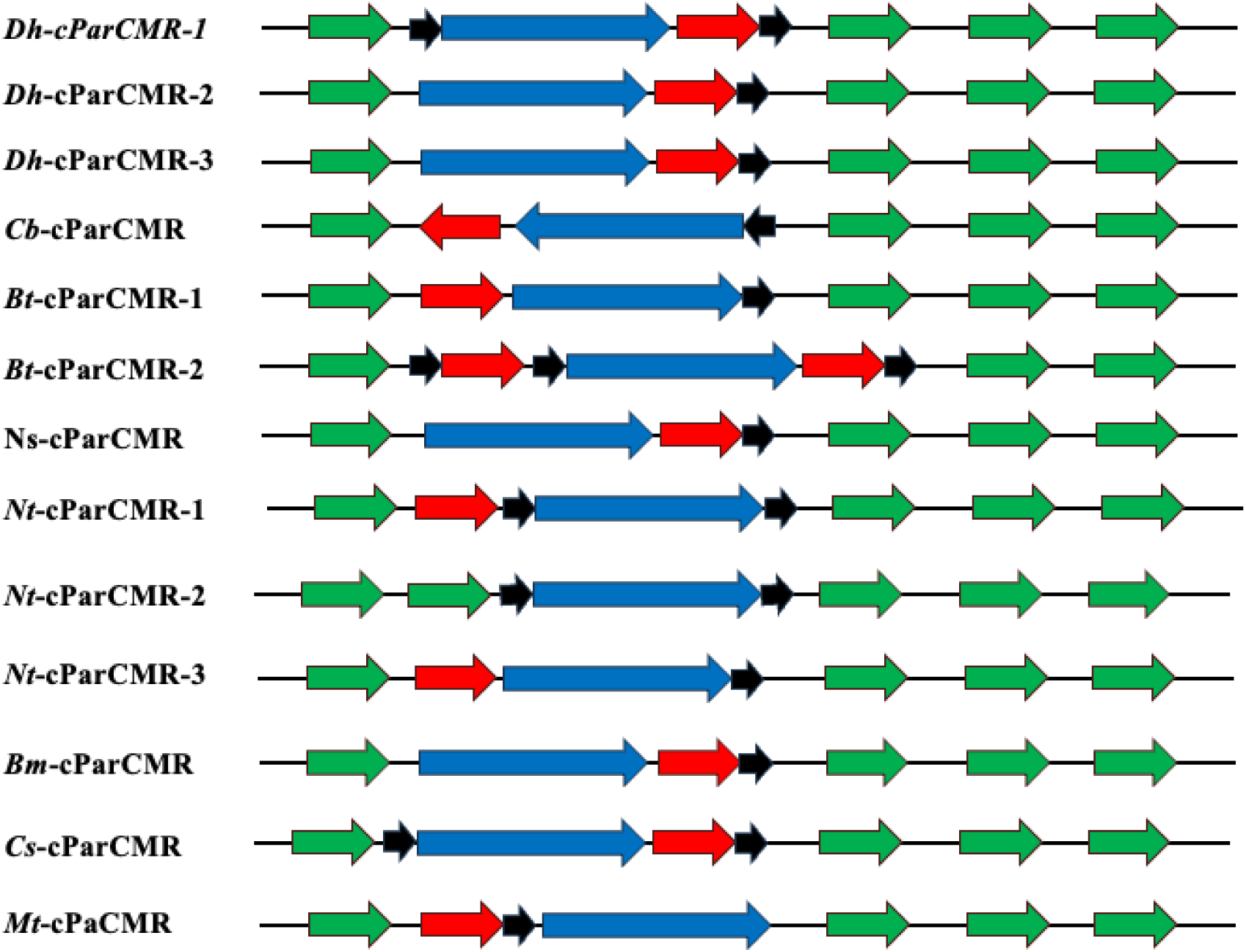
Putative cParCMR cassettes from different bacterial species. Colour coding: cParMs (blue), cParRs (red), c*parC* sequences (black), other unrelated genes (green) (Bengelsdorf et al., 2015; Zhao et al., 2011). Three putative cParCMR systems are encoded on the *Desulfitobacterium hafniense* Y51 chromosome (accession numbers: AP008230.1 or NC_007907.1, 5,727,534 bp): *Dh*-cParCMR-1 (*Dh*-cParM1, WP_011460071.1; *Dh*-cParR1, WP_011460070.1); *Dh*-ParCMR-2 (*Dh*-cParM2, WP_011461902.1; *Dh*-cParR2, WP_041272685.1); and *Dh*-cParCMR-3 (*Dh*-cParM3, WP_011461997.1, *Dh*-cParR3, WP_011461998.1). *Natranaerobius thermophilus* strain JW/NM-WN-LF (accession number: NZ_CP144221.1, 3,137,840 bp) also encodes 3 cParCMR systems - *Nt*-cParCMR-1 (*Nt*-cParM1, WP_148206872.1; *Nt*-cParR1, WP_012449064.1), *Nt*-cParCMR-2 (*Nt*-cParM2, WP_012448769.1) and *Nt*-cParCMR-3 (*Nt*-cParM3, WP_012446843.1; *Nt*-cParR3, WP_012446842.1). This *Natranaerobius thermophilus* strain has two associated plasmids, pNTHE01 and pNTHE02 (accession numbers and sizes: NC_010715.1, 17,207 bp and NC_010724.1, 8,689 bp, respectively). *Bacillus tropicus* strain FDAARGOS_920 (accession number: NZ_CP065739.1, 5,298,747 bp) encodes two putative cParCMR systems *Bt*-cParCMR-1 (*Bt*-cParM1, WP_001968526.1; *Bt*-cParR1, WP_129075283.1) and *Bt*-cParCMR-2 (*Bt*-cParM2, WP_000025611.1; *Bt*-cParR2a, WP_001978111.1; *Bt*-cParR2b, WP_001257752.1). Other bacterial genomes, such as *Moorella thermoacetica* strain 39073-HH (accession number: CP031054.1, 2,645,661 bp), have a single cParCMR cassette (*Mt*-cParM, WP_011391888.1; *Mt*-cParR, WP_053104303.1) and no associated plasmids (Bengelsdorf et al., 2015; Zhao et al., 2011)

### *Dh*-cParCMR-1 cassette

ParM from the *Desulfitobacterium hafniense* Y51 bacterium *Dh*-cParCMR-1 cassette is annotated as a potential chromosome segregating ParM in the NCBI sequence database since the gene resides on the 5,727,534-bp circular chromosome (Nonaka et al., 2006). The protein sequence is highly conserved (percentage identity of 98% to 84%) among five of the six *Desulfitobacterium* species - *metallireducens*, *dichloroeliminans*, *dehalogenans*, *chlororespirans* and *hafniense* (**Table 1**). This high sequence conservation implies an indispensable function in these bacteria. These homologs are variously annotated as “chromosome segregation protein ParM” or “plasmid segregation actin-type ATPase ParM” in NCBI databases. We attempted the *E. coli* expression followed by purification of the *Dh*-cParM1 and *Dh*-cParR1, however only *Dh*-cParM1 produced sufficient protein for characterization.

**Table 1:** *Dh*-cParM1 is highly conserved among the *Desulfitobacterium* species. Sequence identities among five out of the six *Desulfitobacterium species.* cParM was not found by sequence homology searches in *Desulfitobacterium aromaticivoran* genomes.

| Sequence identity (%) | *hafniense* | *chlororespirans* | *dichloroeliminans* | *dehalogenans* | *metallireducens* |
| --- | --- | --- | --- | --- | --- |
| *hafniense* | 100.0 | 98.4 | 94.6 | 92.4 | 84.6 |
| *chlororespirans* | 98.4 | 100.0 | 92.7 | 94.9 | 84.3 |
| *dichloroeliminans* | 94.6 | 92.7 | 100.0 | 93.2 | 85.1 |
| *dehalogenans* | 92.4 | 94.9 | 93.2 | 100.0 | 84.8 |
| *metallireducens* | 84.6 | 84.3 | 85.1 | 84.8 | 100.0 |

### *Dh*-cParM1 filaments shows polymorphism in architecture

*Dh*-cParM1 polymerization induced by ATP was followed by light scattering (**Fig. S2A**). The increase in light scattering was confirmed to be due to filament formation by electron microscopy of negative stained samples. Filaments could be induced by the addition of ATP, GTP and non-hydrolysable nucleotides (**S3A-D**). Sedimentation studies indicated that ADP and GDP have a lower ability to induce polymerization relative to ATP and GTP, respectively, with no discernable increase in sedimented *Dh*-cParM1 on addition of GDP (**Fig. S2B**). On the same EM micrograph under the same buffer conditions, two filament morphologies were observed: single filaments and a tightly coupled double filaments (**Fig. 2A**). Fourier transform analysis of the *Dh*-cParM1 filaments in the micrographs indicated initial helical parameters of twist/rise of 156.0°/23.5 Å for the single filament (**Fig. 2B-C**). Since this morphology was more abundant, using cryo-EM we were able achieve a high-resolution structure reconstruction. Analysis of the 2D classes showed clear secondary structure features highlighting the variations in the filament morphologies (**Fig. 2D**). The single filaments displayed structural features resembling that observed for eukaryotic F-actin and ParM-R1. From a total of 2,845 images and a final selection of 71,060 particles, a 4.0 Å resolution density map of a *Dh*-cParM1 single filament constructed from two parallel strands was reconstructed displaying a helical rise and twist of 24.5 Å and 156.03°, respectively (**Fig. 3A-B**). Its short helical twist is about half that of F-actin or ParM-R1 resulting in a more twisted structure. The two protofilaments are more tightly packed than those of ParM-R1 and pCBH ParMs (Bharat et al., 2015; Koh et al., 2019; Szewczak-Harris and Löwe, 2018).

**Figure 2:**
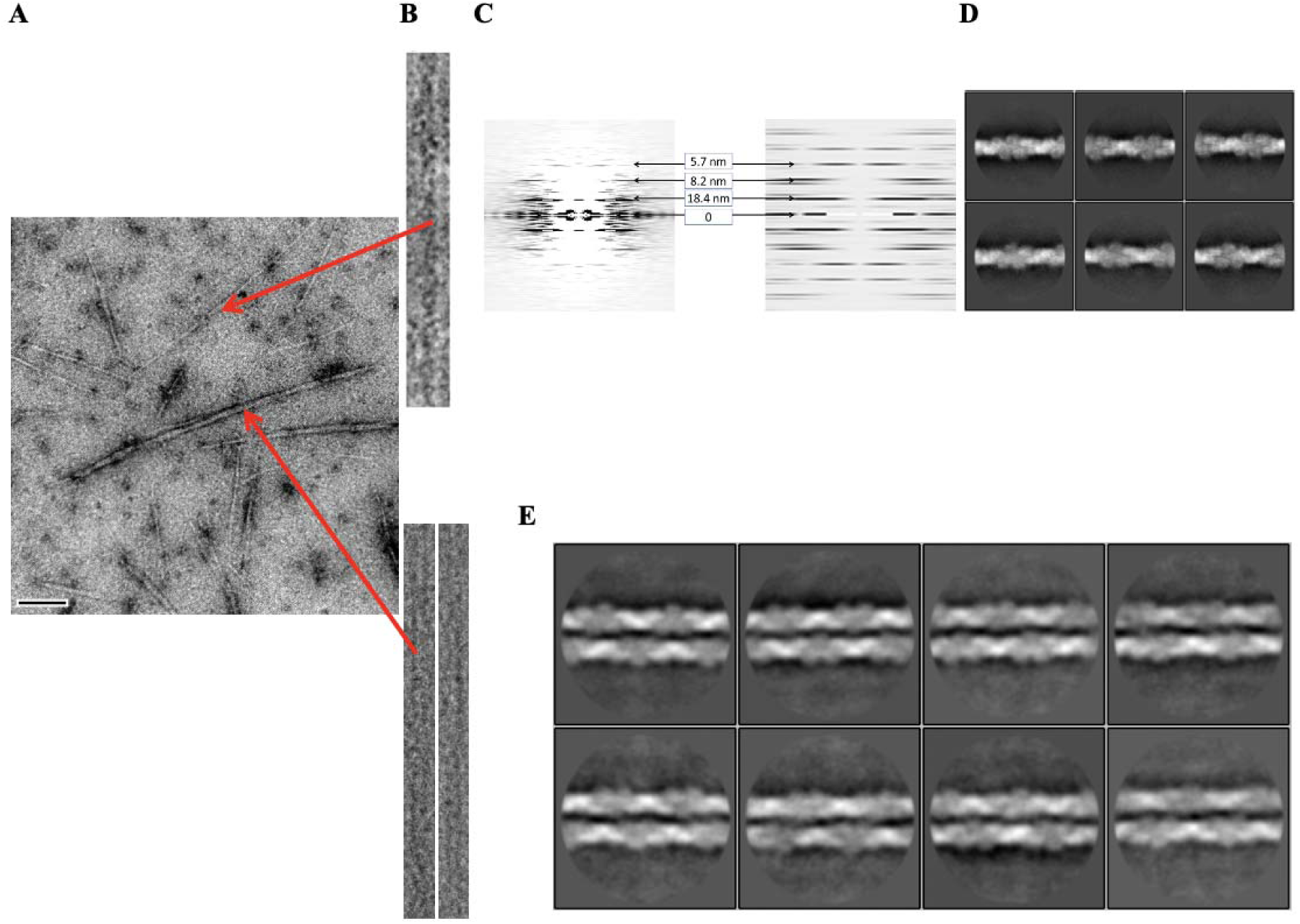
Variations in helical parameters of the two types of *Dh*-cParM1 filaments. **A.** Electron micrograph of a negatively stained sample displaying the two types of *Dh*-cParM1 filament morphologies – single and coupled filaments. **B.** Extracted and straightened single filaments for helical parameters determination. **C.** The averaged diffraction pattern of single filament morphology from negatively stained micrographs (left image) and from reconstructed 3D structure (right image). **D.** 2D classification of *Dh*-cParM1 extracted particles of single filaments from cryoEM micrographs. **E.** 2D class averages of the coupled filament morphology from cryoEM micrographs. Detailed analysis showed these structures comprise of two single filaments lying side by side, which sometimes taper at the ends (**Fig. S3**).

**Figure 3:**
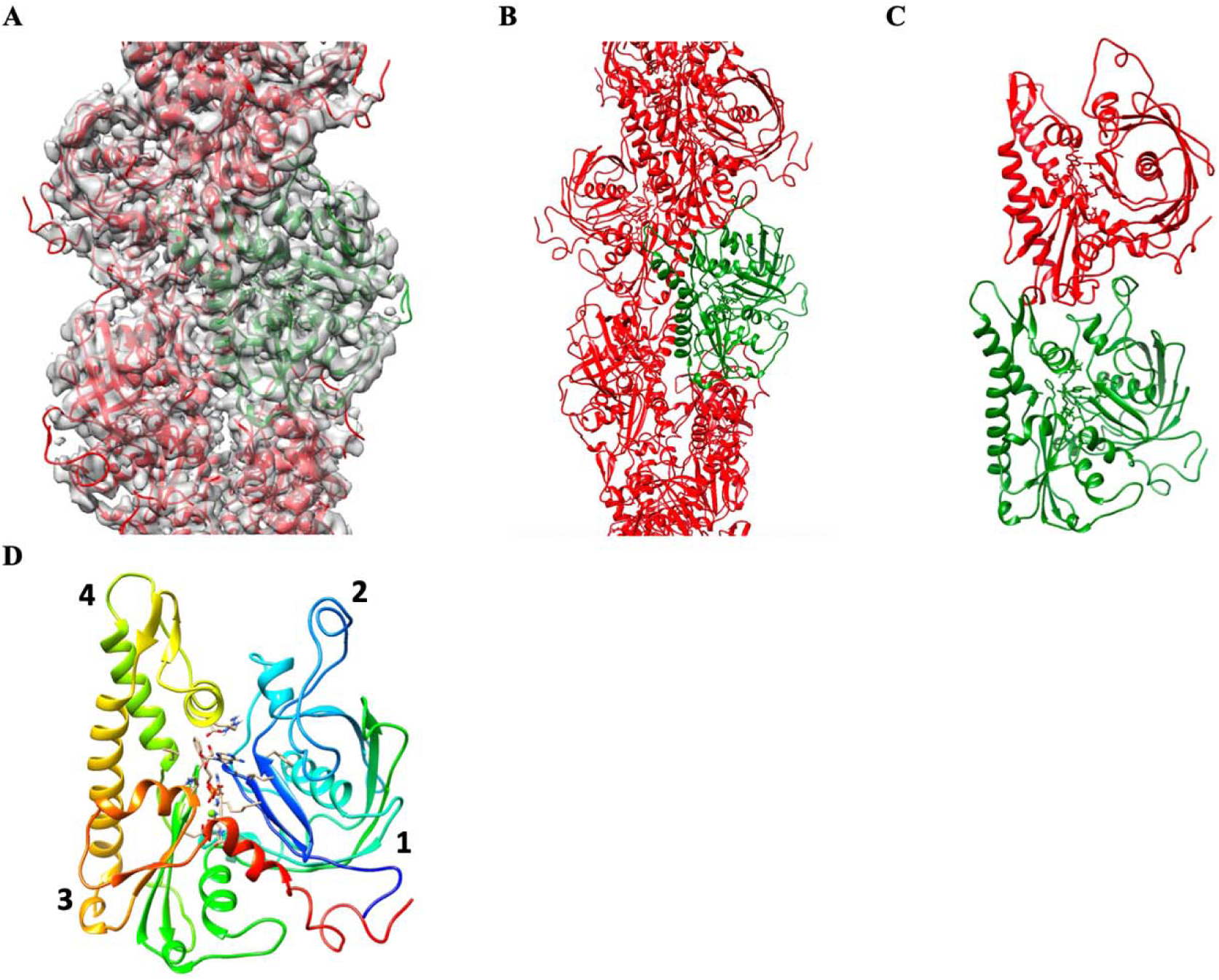
*Dh*-cParM1 shows a double-helical single filament structure. **A.** A protomer fit into the 4 Å resolution density map of *Dh*-cParM1 single filament. **B.** A short segment of the atomic structure of *Dh*-cParM1. One protomer structure is shown in green, the remainder in red. **C.** A dimer of *Dh*-cParM1 protomer showing the longitudinal contacts **D.** *Dh*-cParM1 monomer displaying its four subdomains similar to actin and other ParMs.

We suspected that the tightly coupled *Dh*-cParM1 double filaments may be produced from the side-by-side interaction between two single filaments (**Fig. 2A, E, S3**). The distances between the two filaments were not uniform throughout the structure, often observed to be tapering at the ends. 2D class averaging in RELION 4.0 (with T = 0.5) (Kimanius et al., 2021), confirmed the coupled filament structures (**Fig. 2E**). ParM-R1 can be induced by crowding agents to form similar structures, known as doublets, which show no supercoiling or twisting (Bharat et al., 2015). The interactions between the two *Dh*-cParM1 filaments of this morphology are very similar to those of ParM-R1 doublets. They both showed no supercoiling or twisting of the double filaments. *Dh*-cParM1 spontaneously forms these two filament morphologies without the need for crowding agents. *Dh*-cParM1’s ability to form polymorphs of this nature is a unique feature among ParMs, which may have implications for its function.

### *Dh*-cParM1 protomer and atomic filament structure

We employed AlphaFold2 models as starting models to interpret the cryoEM density maps. Refining these models against the cryoEM density maps, protomer structures from the monomer and filament characterized by good statistics were generated (**Fig. 3A-D, Table S4**). As observed for other ParM protomers, the analogous subdomain 2 region to the DNase I binding loop in actin (Ecken et al., 2016; Kabsch et al., 1990; Wang et al., 2010) was clearly observed in the density as well as the characteristic ParM closed beta-barrel in subdomain 1 (**Fig. 3D**). The hydrophobic residues of subdomain 2 interact with hydrophobic pocket of subdomains 1 and 3 of the upper subunits in a ball and socket in a similar manner as in p*CB*H ParM (Koh et al., 2019) (**Fig. 3C**). The intra-strand arrangements were mainly through lateral longitudinal contacts as seen in other ParMs whereas the inter-strand arrangements are held together by lateral contacts (**Fig. 3B**). The single filament structure is constructed from two parallel, staggered protofilaments.

### *Cb*-cParCMR

*Cb*-cParM (EDT87363.1) and *Cb*-cParR (EDT87283.1) sequences were identified through homology searches using BLAST on a whole genome shotgun sequence of *Clostridium botulinum* Bf (ABDP01000001.1) (**Fig. 1A and Fig. 4**). A putative *parC* sequence containing palindromic sequence repeats was also found immediately upstream of *Cb*-cParMR (**Fig. 5A**). This *Cb*-c*parC* sequence bound to *Cb*-cParR in an electrophoretic mobility shift assay (**Fig. 5B**), indicating that it acts as the *Cb*-cPar*C*. Gene clusters with high similarity to the sequence around *Cb*-cParCMR were found in the genomes of some other *Clostridium* strains and were not found on plasmids (**Figs. 4** and **S4**). Therefore, at the current level of genome sequencing we conclude that *Cb*-cParCMR is encoded on chromosomes from a limited set of *Clostridium* strains.

**Figure 4.**
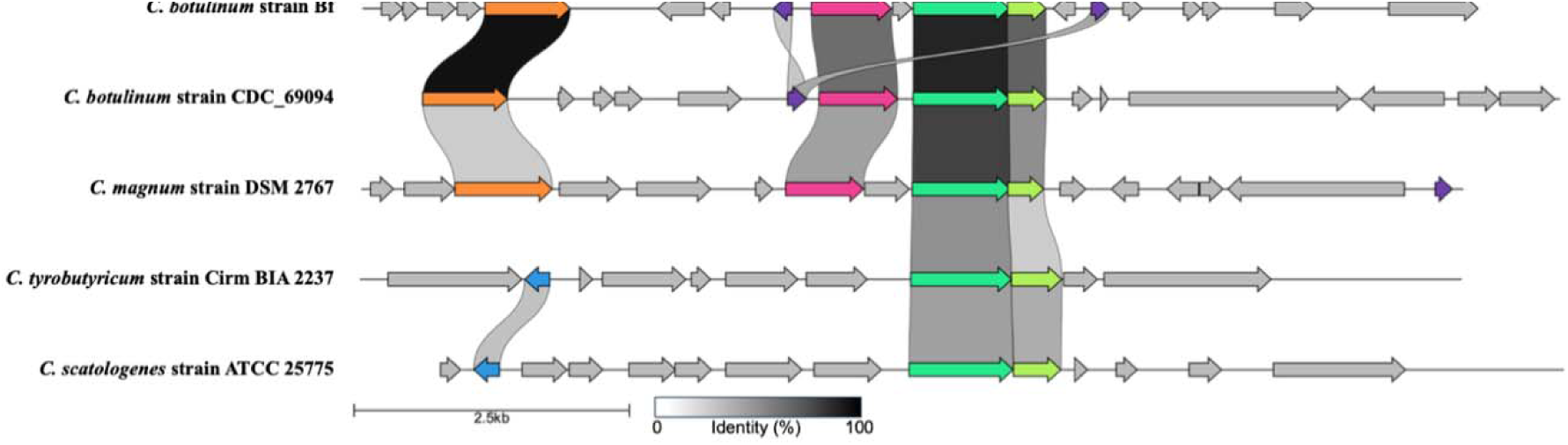
Gene clusters of *Clostridium* species containing cParCMR system. Genes within 5,000 bp of cParM are depicted using clinker & clutermap (Gilchrist and Chooi, 2021). *Clostridium* species containing homologous cParM sequences (identity cutoff = 50 %) are aligned. Genes are depicted as arrows. Conserved genes, cParM (*light green*), cParR (*lime green*), putative replication initiation factor (*magenta*), sporulation-specific N-acetylmuramoyl-L-alanine amidase (*orange*), dihydrodipicolinate reductase (*purple*), and Cro/CI family transcriptional regulator-like protein (*blue*), are colored. Annotations are from GenBank records. c*parC* is not displayed, since a conserved c*parC* sequence was not identified among strains.

**Figure 5:**
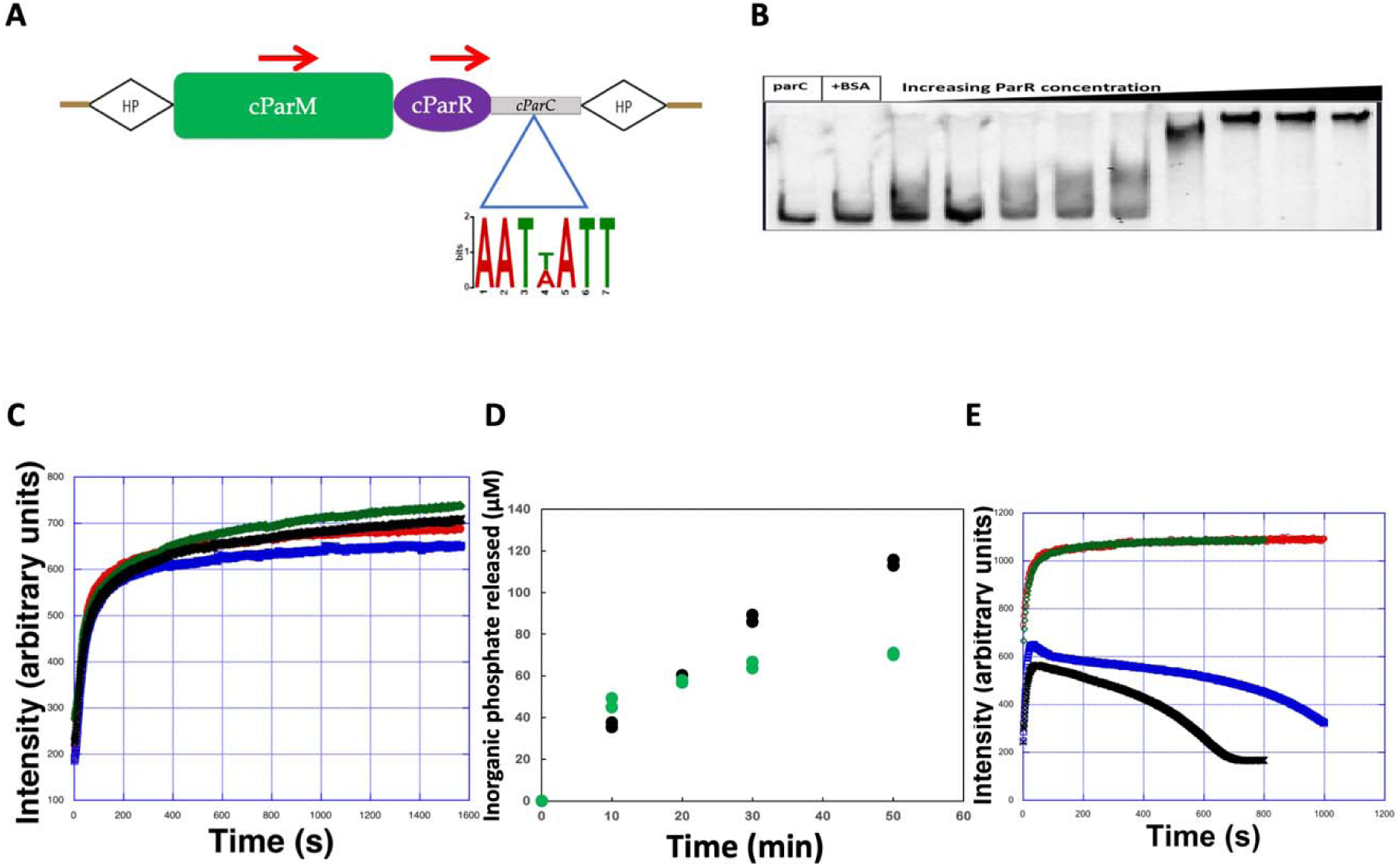
Characterization of the *Cb*-cParCMR system. **A.** cParCMR system present on the genome of *C. botulinum* strain Bf. *Cb*-*cParC* includes three palindromic repeats. HP represents hypothetical proteins on both sides of the cParCMR operon. **B.** Electrophoretic mobility shift assay of *Cb*-*cParC* with 10× to 1000× molar excess of cParR. **C.** Light scattering time courses of *Cb*-cParM polymerization. 10 µM *Cb*-cParM green, ATP; black, GTP; red, ADP; blue, GDP. **D.** Pi release from 21 µM *Cb*-cParM, black with GTP and green with ATP. Two measurements for each nucleotide are superposed. **E.** Light scattering time courses of *Cb*-cParM polymerization in the presence of *Cb*-cParR, blue, with ATP; black, with GTP; Corresponding time courses without *Cb*-cParR, red, with ATP; green, with GTP.

### Polymerization assay and cParR-cParM interactions

*Cb*-cParM polymerized with GTP, ATP, GDP and ADP at similar rates as judged by an increase in light scattering over time (**Fig. 5C**). This is in stark contrast to *Dh*-cParM1 for which polymerization in the presence of ADP and GDP was largely reduced compared to ATP and GTP (**Fig. S2B**). After polymerization, *Cb*-cParM remained as a filament without undergoing bulk depolymerization judged by constant light scattering intensity (**Fig. 5C**), unlike previously studied ParMs (Jiang et al., 2016; Koh et al., 2019) and *Dh*-cParM1 which depolymerized slowly after polymerization (**Fig. S2A**). Supernatant *Cb*-cParM concentrations in a sedimentation assay indicated that the critical concentrations for polymerization were similar in the presence of the different nucleotides, 2.3 ± 0.1 µM, 2.8 ± 0.1 µM, 1.2 ± 0.1 µM, and 3.6 ± 0.4 µM with ATP, ADP, GTP, and GDP, respectively (**Fig. S5**). The critical concentration dependence of *Cb*-cParM on the nucleotide state is considerably smaller than that of actin (Fujiwara et al., 2007) or *E. coli* ParM-R1 (Garner et al., 2004). Continuous phosphate release beyond the concentration of the protein (21 μM *Cb*-cParM) was observed (**Fig. 5D**), suggesting subunit exchange from the ends of the filament after the initial polymerization, consistent with a process such as treadmilling (Narita, 2011; Pollard and Borisy, 2003; Wegner, 1976).

In the presence of *Cb*-cParR, initial polymerization of *Cb*-cParM proceeded normally (**Fig. 5E**) (Garner et al., 2007; Gayathri et al., 2012). However, after the initial polymerization phase, *Cb*-cParR destabilized the *Cb*-cParM filaments and depolymerization occurred, indicating that *Cb*-cParR acts as a depolymerization factor (**Fig. 5E**). A sedimentation assay also indicated the destabilization of the *Cb*-cParM filament by *Cb*-cParR after 30 min incubation (**Fig. S6**).

### CryoEM imaging

We recorded CryoEM images of the *Cb*-cParM filaments after polymerization with ATP or GTP for 30 minutes. Filaments with similar diameters were observed under both conditions (**Figs. 6A-B**). However, 2D classification showed substantially different averaged images (**Figs. 6D-E**). The averaged images of *Cb*-cParM, polymerized with ATP, revealed obvious wide and narrow regions along the filament (**Fig. 6D**), indicating a possible strand cross over. The 2D classification images of *Cb*-cParM polymerized with GTP appeared to show a mixture of two states (**Fig. 6E**). One state was similar to *Cb*-cParM polymerized with ATP, and the other distinct. We tried polymerization with GTP for a shorter incubation time (1 min), which yielded the latter of the mixed states. The averaged images from the short incubation time, showed a more uniform diameter along each filament (**Fig. 6F**).

**Figure 6:**
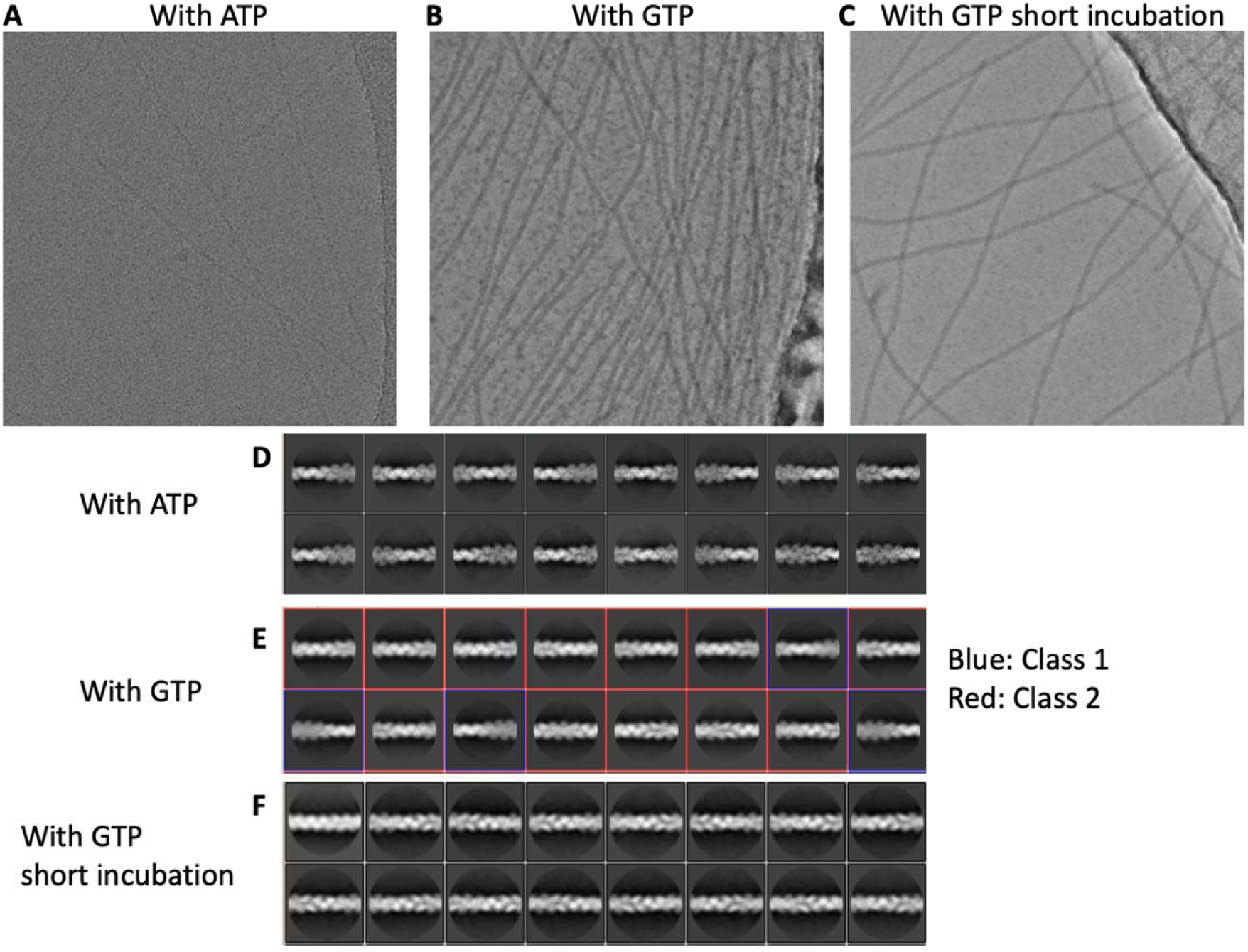
Cryo EM images of *Cb*-cParM filaments. **A, B, C.** Cryo EM images of *Cb*-cParM filament formed with ATP, with GTP, or with GTP and short incubation time, respectively. **D, E, F.** Averaged images after a 2D classification. The images with GTP were classified into two classes: class 1, similar to those with ATP; and class 2, similar to those with GTP and short incubation time.

Four density maps were reconstructed for the three polymerization conditions (**Fig. 7** and **Table S5**). One map corresponds to the first condition (ATP, 30 min, **Fig. 7A**), at 3.9 Å resolution. Two maps were reconstructed from the second condition (GTP, 30 min), corresponding to the two groups in the 2D classification, class 1 and class 2 (**Fig. 6E** and **7B, C**) and one map from the short GTP incubation (1 min, **Fig. 7D**). The map from class 1 at 3.5 Å resolution (GTP, 30 min, **Fig. 7B**) was very similar to that from the first condition (ATP, 30 min, **Fig. 7A**). The two models from the two maps (ATP 30 min, and class 1 with GTP 30 min), were identical up to the reliable resolution of the data, despite the difference in the bound nucleotides. In both maps, density due to gamma phosphate or inorganic phosphate around the nucleotide was not observed, indicating that the bound nucleotides had been hydrolyzed to ADP and GDP, respectively, and the phosphate released. The maps from class 2 with GTP (30 min, **Fig. 7C**) and the third condition (GTP, 1 min, **Fig. 7D**) were similar to each other, although the resolution was limited, suggesting that they were in the state before the phosphate release. For all the 3D reconstructions, the rotation in one strand (twist/rise of ∼48°/49 Å) was very similar to that of *Dh*-cParM1.

**Figure 7:**
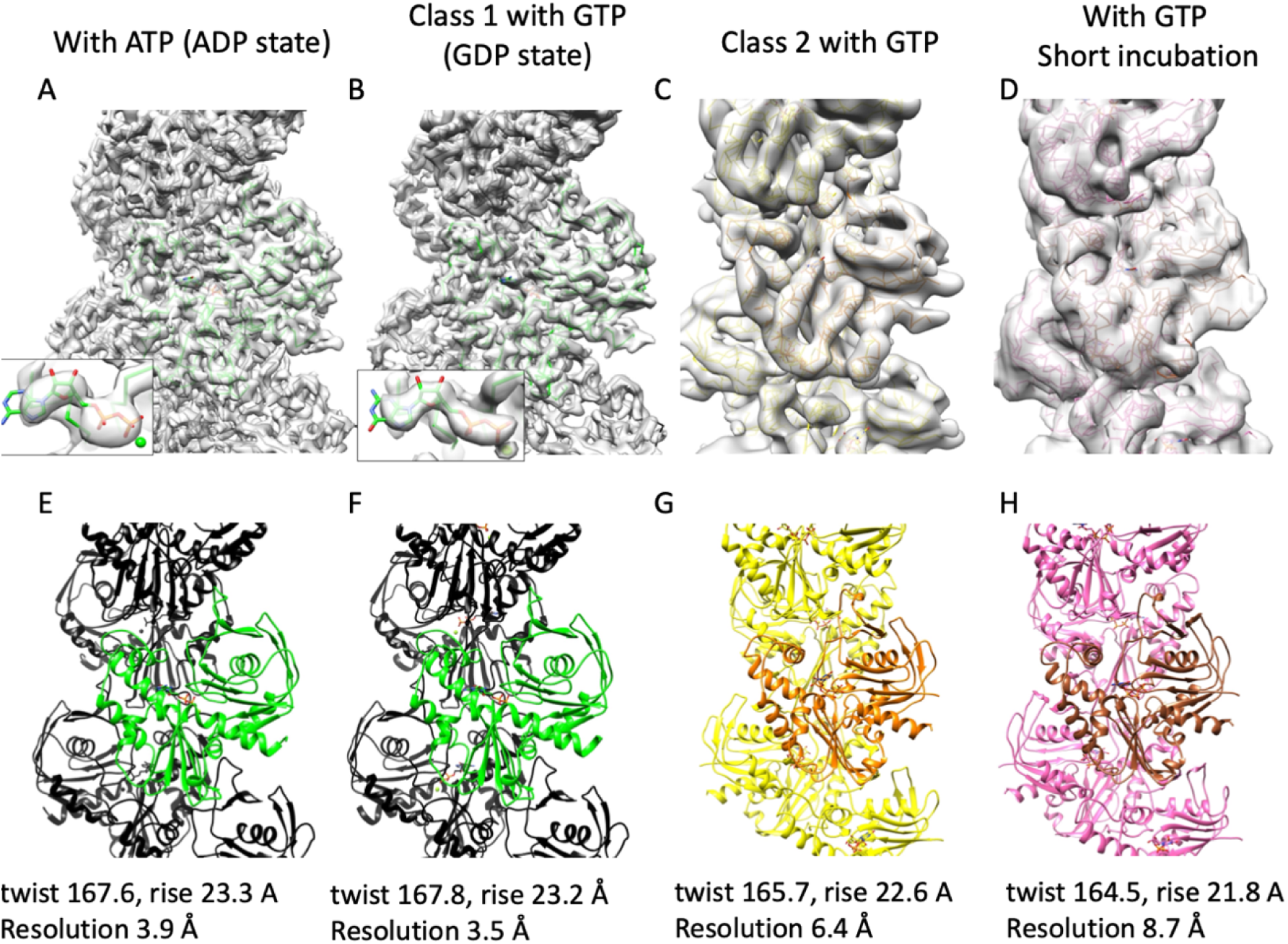
Maps and *Cb*-cParM structural states with bound nucleotides. **A, E.** with ATP. B, **F**. Class 1 with GTP. **C, G**. Class 2 with GTP. **D, H.** with GTP and short incubation time. The structures with ATP (A and E) were almost identical to that of class 1 with GTP (B and F). The gamma phosphate could not be observed in A and B (insets), showing that the binding nucleotides were ADP and GDP, respectively. The model for class 2 with GTP (C and G) and the model with GTP and a short incubation time (D and H) were similar to each other.

### Crystal structure, rigid bodies and domain movement

We successfully obtained a crystal structure of the apo form of a mutant of *Cb*-cParM, without bound nucleotide (**Fig. 8A** and **Table S6**), in which the nucleotide-binding cleft was wide open. This mutant *Cb*-cParM was designed to prevent filament assembly via substitution of three residues in the protomer subunit:subunit interface, R204D, K230D and N234D. We compared the high-resolution model built into the cryoEM map of the GDP state filament (**Fig. 8B**) with the crystal structure of the monomer without nucleotide. We identified two regions that remained almost identical in the large structural change between the models, like those in actin (Oda et al., 2019; Tanaka et al., 2018). We named these regions, the inner domain (ID) rigid body and the outer domain (OD) rigid body (Figs. 8A and **B**), after the actin rigid body nomenclature. In the GDP state, there were no direct interactions observed via hydrogen bonds or salt bridges (**Fig. 8C**), instead the GDP phosphates connected the two rigid bodies (**Fig. 8C**). This explains why the crystal structure without nucleotides adopts wide open state, due to the lack of interactions between ID and OD. The nucleotide binding cleft was slightly more in the open conformation in the models for the class 2 with GTP and short incubation time with GTP, relative to the GDP state (**Fig. 8D**). In addition, the guanine moiety of the GDP did not form contacts or hydrogen bonds with the protein, explaining why *Cb*-cParM can utilize both GTP and ATP. The density of guanine and adenine moieties was relatively weak, indicating the flexibility of these parts due to their lack of interaction with the protein (Figs. 7A and **B**).

**Figure 8:**
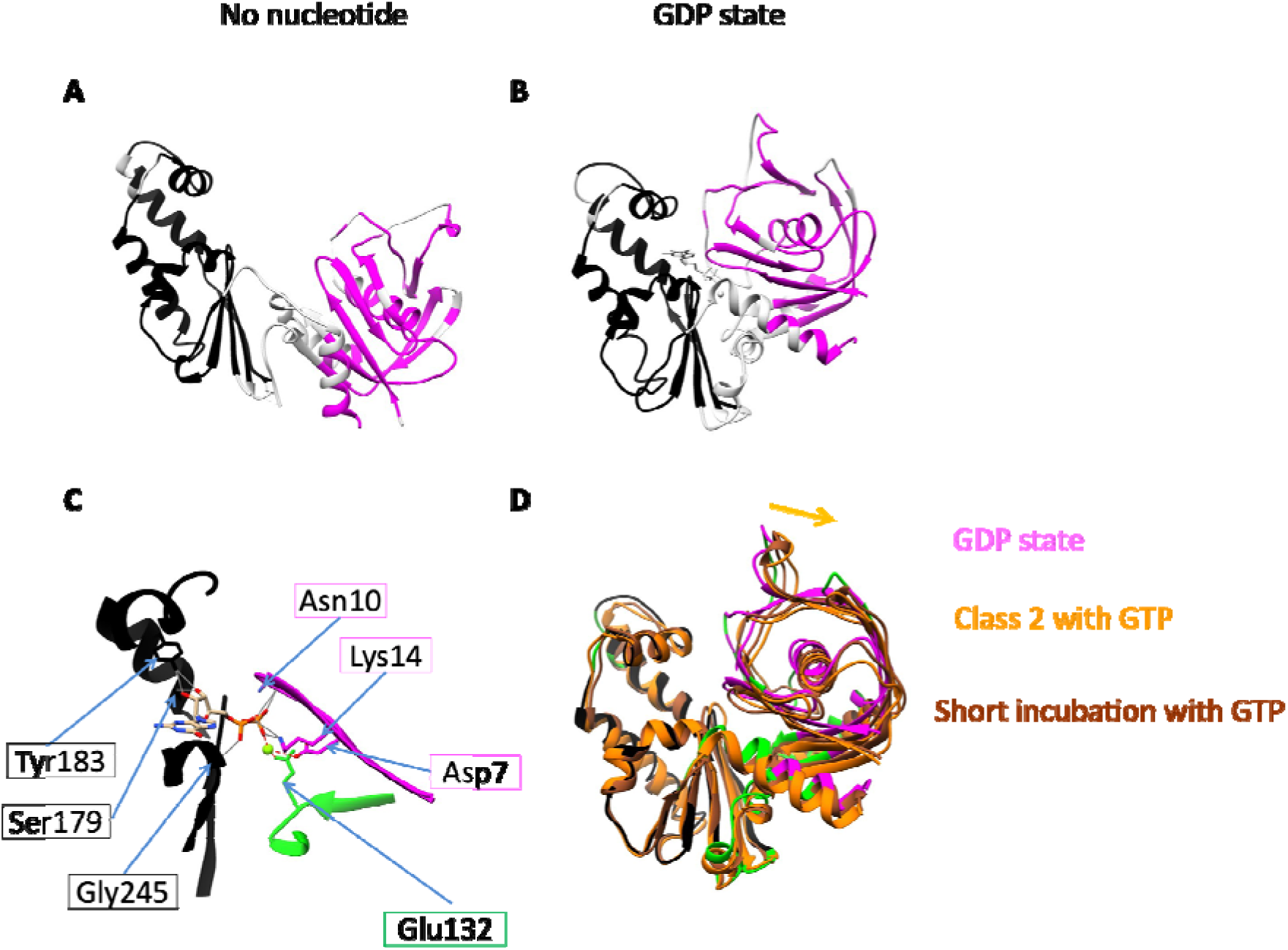
Identification of rigid bodies in *Cb*-cParM. **A.** The crystal structure without nucleotide. **B.** Cryo EM structure with GDP (Fig. 3B). Two rigid bodies were identified by comparing these structures (ID rigid body, black and OD rigid body, magenta). **C.** The bound GDP and Mg^2+^ connected the two rigid bodies (ID rigid body in black, OD rigid body in magenta and the rest of the protein in green). Possible hydrogen bonds corresponding to GDP were determined using UCSF chimera (Pettersen et al., 2004) and presented as gray lines and possible salt bridges with the Mg^2+^ presented by red dotted lines, although additional hydrogen bonds via water molecules may exist. **D.** Models for the class 2 state with GTP (orange) and the short incubation time with GTP (brown) were aligned by the ID rigid body and superposed on the model with GDP (green, black, and magenta). The nucleotide binding cleft was more open in the models for the class 2 with GTP and short incubation time with GTP relative to the GDP state. The orange arrow indicates the direction of domain movement in the open conformation.

### Filament structure

The *Cb*-cParM filament reconstruction with ATP (30 min) was indistinguishable to that of class 1 with GTP (30 min). Therefore, we compared the three remaining filament models: (i) class 1 with GTP (30 min, GDP state), (ii) class 2 with GTP (30 min), and (iii) short incubation time with GTP (1 min). The intra-strand interactions between subunits were very similar in the three models (Fig. 9A and **B**), except for a slight shift in the position of the adjacent subunit in the GDP state. This difference can be explained by the closure of the cleft in the GDP state, which can push the upper subunit leftwards. Despite the similarity within single stands, a large change was observed between strands in forming the filament structures. When the ID rigid bodies of one subunit of each state were aligned with each other, the positions of opposite strands were significantly different (**Fig. 9C-D**, **Movie S1**), with a shift of ∼2.5 nm (Fig. 9D, E). The GTP (30 min) class 2 state and GTP (1 min) are equivalent in their inter-strand orientations and have less cross-strand contacts than observed in the GDP state (**Fig. 9F-G**), suggesting stabilization of the filament bound to GDP. Thus, there are two dramatically different *Cb*-cParM filament conformations: one comprised of loosely associated strands (likely to be bound to GTP or GDP-Pi); and the second formed by a closer association of the strands bound to GDP (**Fig. 9**).

**Figure 9:**
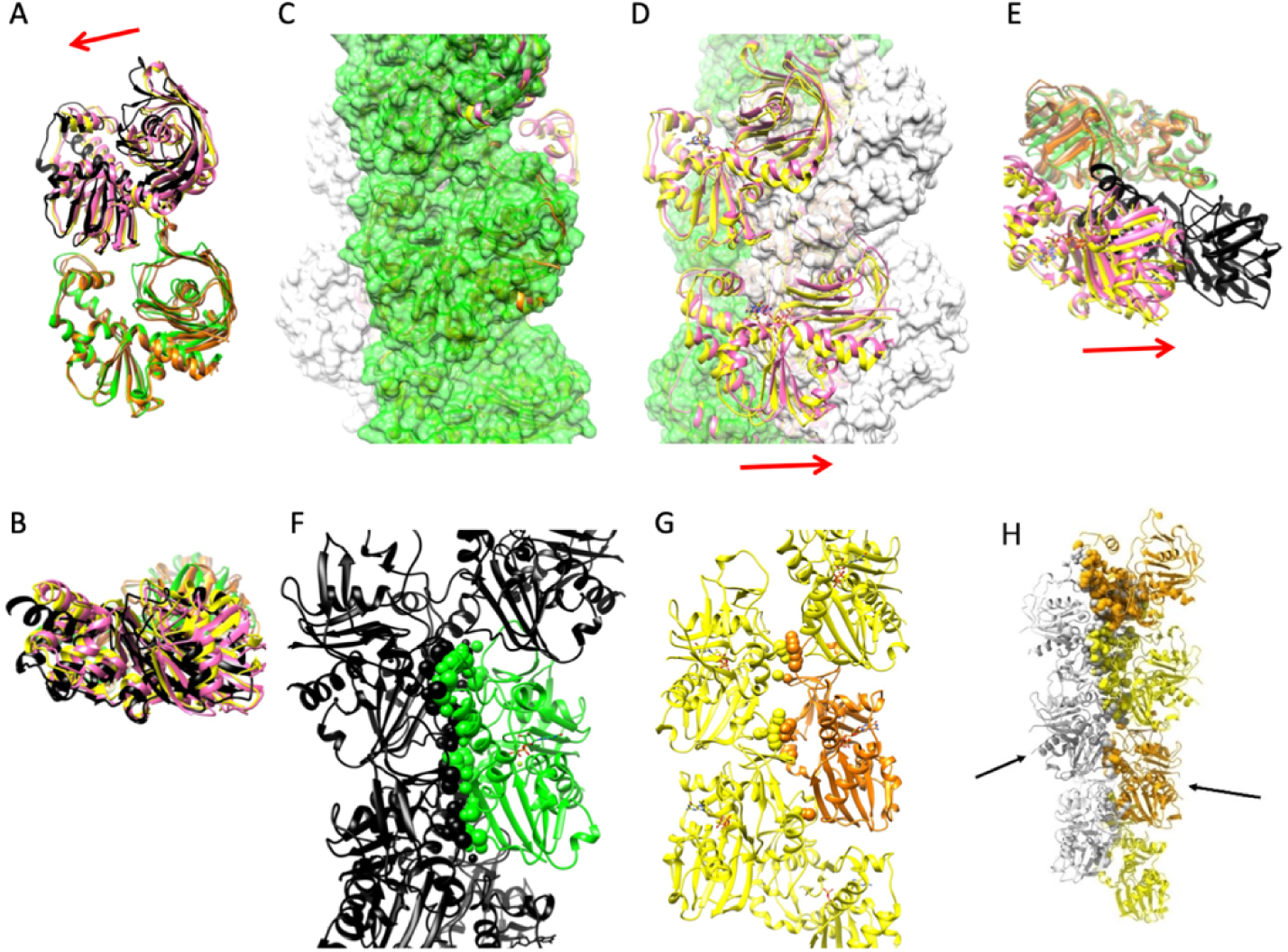
Structural shift in *Cb*-cParM filament strands. **A, B.** Two subunits in one strand (A: front view and B: top view). The ID rigid bodies of the lower subunit were aligned with each other. GDP state: green and black, class 2 with GTP: orange and yellow, short incubation with GTP: brown and pink. The interactions between the subunits appear similar in the three states except for the upper subunit position, which was slightly shifted in the GDP state compared to the other two states. The closure of the cleft in the GDP state may explain this difference because the closure of the cleft can push the upper subunit leftward. A red arrow indicates the direction of the shift of the upper subunit. **C-E.** Relative positions of the two strands. **C.** The model with GDP is presented as a surface model in green and light gray. Ribbon models for the class 2 with GTP (orange and yellow) and the short incubation time with GTP (brown and pink) are superposed. The ID rigid bodies of the center subunits are aligned with each other. **D.** 180 ° rotated view of Fig. 5C. The opposite strand position in the GDP state was completely different from that in the other two states (compare gray surface to cartoon). **E.** Top view of two adjacent subunits in the different strands. GDP state: green and black, class 2 with GTP: orange and yellow, short incubation with GTP: brown and pink. The ID rigid bodies of the lower subunit (green, orange and brown) were aligned with each other. A red arrow indicates the direction of the shift in D and E. **F.** Inter-strand interactions in the *Cb*-cParM filament model with GDP. Atoms from the center molecule (green) in contact with the other strand (black) are presented in space filling representation. 176 contacts are observed. **G.** Inter-strand interactions in the model for the class 2 with GTP. 36 contacts were observed. **H.** The strands of the model with GDP (light gray) were replaced by the strands of the model for the class 2 with GTP (orange and yellow). The strands were aligned by the molecules indicated by black arrows. The clashing atoms were presented in space filling representation. The contacts and clashes were identified by ChimeraX (Goddard et al., 2018).

## Discussion

Orthologs of plasmid-encoded ParABS systems have been identified on bacterial chromosomes (Mohl and Gober, 1997; Ptacin et al., 2010). The molecular mechanisms by which these partitioning genes segregate plasmids and chromosomes have been explored in several bacteria species. Unlike the ParABS systems which function both as plasmid and chromosome segregation systems, only a plasmid partitioning function for ParCMR systems has been established. Here, our findings demonstrate that the biochemical, biophysical, and structural properties of ParCMR homologs located on bacterial chromosomes are similar to those of plasmid-borne ParCMR systems.

Biochemically, both *Dh*- and *Cb*-cParMs polymerize to form filaments in a nucleotide-dependent manner (**Fig. S2, S3, 5C-E, 6A-C**). Furthermore, protomers of the characterized cParMs (**Fig 3D, 8D**) clearly possess the characteristic ParM-closed beta-barrel domain, (van den Ent et al., 2002) comprised of a three-stranded β-sheet wrapped around a helix, in a similar fashion as ParM-R1 (van den Ent et al., 2001), pSK41 plasmid ParM (Popp et al., 2010), BtParM (Jiang et al., 2016), AlfA (Szewczak-Harris and Löwe, 2018) and p*CB*H ParM (Koh et al., 2019). AlphaFold2-generated models (**Fig. S1**) indicate that the wider set of genome-borne ParMs also exhibit the beta-barrel subdomain. This demonstrates that the ParM systems borne on genomes bear resemblance and polymerization activities similar to plasmid borne ParMs. Thus, cParMs likely have functions other than plasmid segregation, which are yet to be explored.

*Cb*-cParM polymerizes with ADP or GDP (**Fig. 5C**), whereas *Dh*-cParM is similar in this respect to other characterized ParMs (Garner et al., 2004; Jiang et al., 2016; Koh et al., 2019). *Dh*-cParM polymerizes with ATP or GTP and is destabilized with ADP or GDP (**Fig. S2B**). It is likely that for *Dh*-cParM, polymerization with GDP was either short-lived or totally absent. The *Cb*-cParM filaments, polymerized with either di-phosphate nucleotide, were stable (**Fig. S5**). Phosphate release from *Cb*-cParM polymerized from ATP or GTP did not cause filament instability, as observed for other ParM systems, rather the filament underwent a significant structural change. More inter-strand contacts were observed in the GDP-bound state, providing stability after phosphate release (Fig. 9F and **G**). When the strands in the GDP model were replaced by the class 2 model with GTP, in silico, severe clashes were observed (**Fig. 9H**). Therefore, the small change in the strand structure (**Fig. 9A**) caused by the opening of the nucleotide binding cleft (**Fig. 8D**) in the class 2 model with bound GTP prevents the formation of the close contacts observed in the model with GDP. After phosphate release, the strands adopt the GDP conformation allowing for new inter-strand contacts. Thus, phosphate release results in a large structural change in the filament.

The *Cb*-cParM cryoEM images showed obvious differences in the distribution of the nucleotide states between ATP and GTP at 30 mins after initiation of polymerization. It implies either that phosphate release from the ADP-Pi state is faster than that from GDP-Pi state, or faster nucleotide hydrolysis, shortening time in the states of ADP-Pi and ATP in the nucleotide hydrolysis cycle. This is consistent with the faster phosphate release observed with ATP under steady state conditions following the initial polymerization (**Fig. 5D**, gradient from 10 mins).

We discovered that *Cb*-cParM filaments depolymerized by a different mechanism to filament instability following phosphate release. *Cb*-cParR acts as a depolymerization factor for the aged filament. We speculate that *Cb*-cParR recognizes the large structural change in *Cb*-cParM after phosphate release allowing it to change its role to depolymerization. This mechanism has parallels in the eukaryotic actin system, where cofilin senses the nucleotide status of the actin filament, resulting in depolymerization (Carlier et al., 1997). Since, *Cb*-cParR also binds to *Cb-cparC* it appears that *Cb*-cParR may play dual roles. The DNA-bound *Cb*-cParR may bind to *Cb*-cParM filaments, by analogy to plasmid ParCMR systems, while the free *Cb*-cParR acts as a depolymerizing factor for aged *Cb*-cParM filaments.

These two described bacterial genome-borne ParCMR systems have been proven to be functional and active polymerization systems like those present on segregating plasmids. Additionally, we have discovered many other systems on genomes of bacteria which may, or may not, also possess plasmids (a representative selection is shown in **Fig. 1**). Whereas some cParMs such as *Nt*-cParMs are not conserved on the chromosome of *Natranaerobius* species*, Bt*- and *Mt*-cParMs, like *Dh*-cParM, are highly conserved in *Bacillus* and *Moorella* species, respectively (Table **S1-S3**). The high sequence conservation within these bacterial species especially *Desulfitobacterium* species, together with the striking feature of the absence of plasmids in this bacterium (Nonaka et al., 2006), strengthens the hypothesis of indispensable functions other than plasmid partitioning. We cannot rule out the possibility that the cParMs encoded on genomes contribute to plasmid segregation when plasmids are present. However, in the case of the *Cb*-cParCMR cassette, a functional (*Cb*-cParR binding) *parC* is borne on the genome, implying this cassette most probably is involved in different role than plasmid segregation. Currently, only type I plasmid partitioning ABS systems have been implicated in chromosome segregation of *C. crescentus*, *Vibrio cholerae* and *B. subtilis* (Fogel and Waldor, 2005; Kadoya et al., 2011; Lewis and Errington, 1997; Mohl and Gober, 1997). The identification of genome borne ParCMR homologous systems paves the way for further studies to experimentally explore whether these systems are functional equivalents of chromosome segregating ParABS system, or whether they play other undiscovered roles.

## Materials and Methods

### Bioinformatics analysis and identification of ParCMR operons

A bioinformatics sequence database search (NCBI and Uniprot databases) was performed on genomes from bacteria to identify the presence of the ParCMR loci using the conserved actin superfamily fold, including the nucleotide-binding region of the ParM sequence (Kabsch and Holmes, 1995). ParR and *parC* were found either upstream or downstream of the ParM gene (Adams et al., 2015). Unlike ParMs, which inherit the nucleotide-binding region for easy identification, ParR does not possess such a key motif. *parCs* are usually characterized by short repeat DNA sequences were identified accordingly. Finally, a DELTA-BLAST search was used to locate potential overlooked ParM and ParR sequences.

### Protein Expression and Purification

Constructs of ParM and ParR were gene synthesized and cloned into pSY5 and pSNAP vectors, respectively, encoding an 8-histidine tag followed by a human rhinovirus HRV 3C protease cleavage site. Plasmids were transformed into BL21 (DE3) cells grown to OD_600_ ∼ 0.8, and protein expression was induced with 0.2–1.0 mM IPTG overnight at 15 °C. The cultures were then centrifuged at 4000 × *g* and the cell pellets were resuspended in 50 mM Tris–HCl pH 8.0, 500 mM NaCl, 20 mM imidazole, 5% glycerol, 0.5 mg/mL lysozyme, 0.1 % Triton-X, and protease inhibitor tablets (1 per 2 L culture, Roche, Basel, Switzerland) and lysed via sonication. The cell lysate was then clarified by centrifugation at 30,000 × *g* for 30 min and filtered through a 0.45 µm membrane. The filtered supernatant was loaded onto a HisTrap FF 5 mL (GE Healthcare, Marlborough, MA, USA) pre-equilibrated with 50 mM Tris-HCl (pH 8.0) containing 500 mM NaCl and 20 mM imidazole. Following a washing step, human rhinovirus HRV 3C protease (5 mg/ml) was loaded in the same buffer for cleavage of tagged proteins (12 h at 4 °C). The cleaved protein was then eluted with washing buffer, pooled, concentrated and subjected to size-exclusion chromatography (Superdex 75 pg, GE Healthcare) in 40 mM HEPES pH 7.5, 150 mM KCl, 2 mM MgCl_2_, and 1 mM DTT. Fractions were checked for purity via SDS–PAGE, and the pure fractions were pooled and concentrated to between 5 and 10 mg/mL, as determined by UV absorbance at 280 nm using an estimated A_280_ value calculated using PROTEINCALCULATOR v3.4 (http://protcalc.sourceforge.net).

### Electrophoretic mobility shift assay

The reaction mixture (10 µL) containing 20 nM to 20 µM of ParR in 25 mM HEPES-HCl (pH 7.5), 300 mM KCl, 1 mM MgCl_2_, 0.5 mM DTT, 1 mg/ml bovine serum albumin, 0.1 µg/µl sonicated salmon sperm DNA, and 5 % glycerol was mixed at 25 °C for 10 min, followed by the addition of 20 nM 5’-FAM-labelled *parC* DNA fragments and further incubation for 20 min. The polyacrylamide gels were prerun at 150 V for 1 h. After incubation, reactions were analyzed by electrophoresis on a 1 x TBE (pH 7.5), 4% polyacrylamide gel in 1 x TBE running buffer (0.89 M Tris-base, 0.89 M boric acid, 0.02 M EDTA, pH 8.3) at 150 V for 1 h. Gels were scanned using a Pharos FX Plus Molecular Imager (Bio-Rad, Hercules, CA, USA) with an attached external laser.

### Polymerization assays

Assembly and disassembly of ParMs at 24 °C was followed by light scattering at 90 ° using a Perkin Elmer LS 55 spectrometer for extended time measurements (initial delay time due to mixing by hand ∼ 10-50 s) or a BioLogic stopped-flow machine to observe the early polymerization phase (initial delay time ∼ 3 ms), monitored at 600 nm. Assembly was initiated by addition of 2 mM nucleotide at 24 °C in 20 mM Hepes pH 7.5, 350 mM KCl, 2 mM MgCl_2_ (*Cb*-cParM) and 40 mM HEPES, pH 7.5, 400 mM KCl, 1 mM MgCl_2_ (*Dh*-cParM).

### Pi release assay

*Cb*-cParM protein (21 µM) was mixed with the appropriate nucleotide in the same buffer as the cryoEM conditions and incubated according to the time course. The reaction was stopped by adding equal volume of ice-chilled 0.4 M perchloric acid (PCA). The reaction mixture was then centrifuged at 1,700 x g for 1 minute. Equal volumes of supernatant and Malachite Green reagent were mixed and incubated at 25 °C for 25 minutes (Kodama et al., 1986). Absorbance was recorded at 650 nm with a Hitachi U-3010 spectrophotometer.

### Sedimentation Assay

The polymerization of *Dh*-cParM (20 µM) with different nucleotides was monitored after addition of 5 mM nucleotide (ATP, ADP, GTP, or GDP) in 40 mM HEPES (pH 7.5), 300 mM KCl, 2DmM MgCl_2_, and 0.5 mM DTT buffer at 24 °C for 30 min. Samples were centrifuged at 279,000 × *g* for 20 min. Pellets were resuspended in the same volume of buffer as reaction mixture. Supernatant and pellet fractions were analyzed through SDS-PAGE.

To investigate the critical concentrations for polymerization, polymerization of different concentrations of *Cb*-cParM (4–15 µM) was initiated by the addition of 5 mM nucleotide (ATP, ADP, GTP, or GDP) in 40 mM HEPES (pH 7.5), 300 mM KCl, 2DmM MgCl_2_, and 0.5 mM DTT at 24 °C for 30 min. Samples were centrifuged at 279,000 × *g* for 20 min and pellets were resuspended in the same volume as the reaction. Concentrations of *Cb*-cParM in the supernatant were estimated via SDS–PAGE, and gel images were analyzed using ImageJ software. The concentrations were not dependent on the total concentration of *Cb*-cParM, indicating that they reflected the critical concentration for each nucleotide state. Therefore, the concentrations of the supernatants from different total ParM concentrations were averaged.

To investigate the effects of *Cb*-cParR on *Cb*-cParM, polymerization of *Cb*-cParM (20 µM) with and without *Cb*-cParR (20 µM) was initiated by the addition of 5 mM nucleotide (ATP, ADP, GTP, and GDP) in 40 mM HEPES (pH 7.5), 300 mM KCl, 2DmM MgCl_2_, and 0.5 mM DTT at 24 °C for 30 min. Samples were centrifuged at 279,000 × *g* for 20 min and pellets were resuspended in the same volume as the reaction and the concentrations of *Cb*-cParM in the supernatant were estimated via SDS–PAGE.

### Crystallography

*The Cb*-cParM mutant was constructed with three mutations (R204D, K230D and N234D) to prevent polymerization and allow for crystallization. Purified protein was subjected to crystallization trials by mixing and incubating 5 mg/ml *Cb*-cParM mutant and 1 mM AMPPNP on ice for 1 hour. Via the hanging drop vapour diffusion method crystals were grown in 0.5 µl of protein/AMPPNP and 1 µl of mother liquor (0.2 M ammonium chloride, 22% (w/v) PEG 3350) at 288DK. X-ray diffraction data were collected on a RAYONIX MX-300 HS CCD detector on beamline TPS 05A (NSRRC, Taiwan, ROC) controlled by BLU-ICE (version 5.1) at λD=D1.0DÅ. Indexing, scaling, and merging of data was performed using HKL2000 (version 715) (Otwinowski and Minor, 1997). Molecular replacement using the protomer built into the 3.5 Å cryoEM density map was carried out in the Phaser (Adams et al., 2011) after splitting the structure into the two domains. Model building was carried out in Coot (Emsley and Cowtan, 2004) and refinement in PHENIX (Adams et al., 2011). Data collection and final refinement statistics are summarized in Table S1. Although *Cb*-cParM was crystallized in the presence of AMPPNP, the resultant structure did not contain nucleotide.

### Cryo-electron microscopy of *Dh*-cParM

ParM protein (20 µM) was polymerized with 5 mM ATP in 40 mM HEPES, pH 7.5, 400 mM KCl, 1 mM MgCl_2_ buffer at room temperature for 20-30 minutes. The mixture (2.5 µl) was applied to previously glow discharged R 1.2/1.3 Molybdenum 200 grids with a holey carbon support film (Quantifoil). Grids were quickly transferred to an EM GP Leica for blotting for 1.5-2.5 seconds at 90% humidity and flash frozen in liquid ethane cooled by liquid nitrogen. Grids were stored under liquid temperatures and screened on a Tecnai G2 Polara (FEI, Nagoya University) operated at 300 kV at minimal dose system. Final cryoEM data were collected on a Titan Krios (FEI, Osaka University) equipped with FEG operated at 300 kV and a minimal dose system. Imaging was done using the EPU software (FEI) attached to the Titan Krios. Images were recorded at nominal magnification of 75,000, objective aperture of 100 µm, actual defocus range between −1.5 to −2.5 µm with a dose rate of 45De−/Å^2^/s and exposure time of 1Dsecond with three image acquisitions per hole. Images were recorded using a Falcon II detector (FEI) at a pixel size of 1.1 Å/pixel and a frame rate of 17Dframes/s. The first two frames were discarded and successive seven frames saved for image processing.

### Image processing of *Dh*-cParM

About 2200 to 2500 images were collected from different microscope sessions and processed using RELION 2.0/3.0/4.0 (He and Scheres, 2017; Kimanius et al., 2021; Zivanov et al., 2018). Frames were motion corrected with MotionCorr (Li et al., 2013) and CTF estimation performed with Gctf (Zhang, 2016). Micrographs with good observed CTF were selected for further processing. Using e2helixboxer, filaments were manually picked and extracted at a box size of 400 x 400 pixels in RELION 2.0/3.0/4.0. Particles from 2D classes displaying clear secondary structure elements were selected. Initial 3D reference models were prepared using RELION toolbox kit cylinder or reconstructed 3D model from negative stain images using EOS software (Yasunaga and Wakabayashi, 1996). A 3D refinement was performed with a low pass filter of 40 Å and helical symmetry search without solvent flattening. Converged helical symmetry was obtained and additional 3D refinement performed with determined helical parameters with solvent flattening. Particle polishing, movie refinement without CTF refinement and final 3D refinement were performed using the previous 3D refinement as initial model producing a 4.0 Å map of *Dh*-cParM converged to a rise of 24.5 Å and twist of 156.03°. With post processing using solvent flattening and a soft mask, final resolutions were attained for the ParM filaments. Handedness was determined by the handedness of the alpha helices observed in a single strand of the high resolution filament structure.

### *Cb*-cParM Cryo-electron microscopy

*Cb*-cParM (0.7 mg/mL) was polymerized in 20 mM HEPES-HCl pH 7.5 containing 250 mM KCl, 1.7 mM MgCl_2_, 3 mM GTP or ATP. The mixed solution was incubated for 30 min or 1 min at 25°C. R1.2/1.3 Mo400 grids (Quantifoil, Jena, Germany) were glow discharged and used within an hour. The reaction mixture (2.5 µL) was applied on the glow discharged grids, blotted on the EM GP (Leica, Wetzlar, Germany) and vitrified by plunging in liquid ethane cooled by liquid nitrogen. Frozen grids were kept under liquid nitrogen for no more than 1 week before imaging. To screen for optimum conditions for cryoEM imaging, the grids were manually observed in a Tecnai G2 Polara (FEI, Hillsboro, OR, USA) cryo transmission electron microscope (at Nagoya University) equipped with a field emission gun operated at 300 kV and a minimal dose system. Images were captured at a nominal magnification of ×115,000 with an underfocus ranging from 1.5 to 3.5 μm and by subjecting the sample to a 2 s exposure time corresponding to an electron dose of ∼30 electrons per Å^2^. Images were recorded on a GATAN US4000 CCD camera using an energy filter operated between 10 and 15 eV, with each pixel representing 1.8 Å at the specimen level at exposure settings. Samples were imaged using a Titan Krios microscope operated at 300 kV installed with EPU software (Thermo Fisher, Waltham, MA, USA) at Osaka University. The imaging parameters were nominal magnification of 75,000 and 91,000, actual defocus 1.0–3.0 μm, dose rate 45 e^−^/Å^2^/s, exposure time 1 s, and three image acquisitions per hole. The images with ATP were recorded with a Falcon II/III detector (FEI) (Thermo Fisher) at a pixel size of 0.87 Å/pixel with an objective aperture 100 µm, while the images with GTP and 30 min incubation time were recorded with a Falcon II/III detector (FEI) (Thermo Fisher) at a pixel size of 0.87 Å/pixel with an objective aperture 100 µm. The images with GTP and 1 min incubation were recorded with a Falcon III (Thermo Fisher) at a pixel size of 0.87 Å/pixel with a phase plate.

### *Cb*-cParM image processing

From *Cb*-cParM samples with ATP, 2,778 images were collected. Image processing was performed using RELION 3.1 (He and Scheres, 2017; Scheres, 2012) software. After motion correction and contrast transfer function (CTF) estimation with CTFFIND-4.1 (Rohou and Grigorieff, 2015), images were selected for further image processing. Filaments were manually picked with e2helixboxer, after which particles were extracted at a box size of 384 × 384 or 400 x 400 pixels. After 2D classification, 36,762 and 71,060 particles respectively were selected. The initial 3D reference was prepared using conventional helical reconstruction using EOS (Yasunaga and Wakabayashi, 1996). Helical symmetry converged to 167.6° twist/22.3 Å rise along the left-handed helix, and the resolution reached 3.9 Å. With *Cb*-cParM GTP and 30 min incubation time, 1,398 images were collected. After motion correction and CTF estimation, 152,490 particles were extracted. We categorized 2D averaged images into two by visual inspection. The first category (class 1) contained 40,599 particles and the helical symmetry converged to 167.8 ° twist/22.3 Å at 3.5 Å resolution, while the second (class 2) contained 70,754 particles and helical symmetry converged to 165.7° twist/21.7 Å rise at 6.5 Å resolution. For the map with GTP and 1 min incubation time, 2,772 images were collected, and 153,326 particles were used for the final reconstruction at 8.6 Å resolution.

### Model building

The initial atomic model of *Cb*-cParM with GDP was constructed by homology modeling using Rosetta3 with the p*CB*H ParM model (Koh et al., 2019) (6IZV) as a template. The resulting model was iteratively refined using COOT (Emsley et al., 2010), molecular dynamics flexible fitting (MDFF, using ISOLDE (Croll, 2018), an extension of ChimeraX (Goddard et al., 2018), and Phenix (Adams et al., 2010). GDP in the final GDP model was replaced by ADP to give the initial model with ADP, which was then refined using the same procedures. The final GDP model was also used as the initial model for the lower resolution structures (PDBIDs 7X55 and 7X59), which were fitted into the map by MDFF with adaptive distance restraints for the two rigid bodies using ISOLDE (Croll, 2018). The resultant model was refined using COOT and Phenix.

For Dh-cParM1, we used AlphaFold2 software (Senior et al., 2020) to generate initial models. The model was refined with the cryoEM density map using ISOLDE software (Croll, 2018), Rosetta program (Rohl et al., 2004) and subsequently refined using PHENIX real space refine to produce the protomer structures.

### Rigid body search

For *Cb*-cParM, the model with GDP and the crystal structure without nucleotides were aligned with each other to maximize the number of Cα with less than 0.7 Å deviation between the two models. The resultant residues with less than 0.7 Å deviation were considered as the rigid body (Tanaka et al., 2018). Two rigid bodies were identified (**Fig 8**).

## Acknowledgments

This research was supported by JSPS KAKENHI (grant numbers 18H02410 and 21H02440 to AN), JST CREST (JPMJCR19S5 to RCR and AN). This research was also supported by Nanotechnology Platform Program of the Ministry of Education, Culture, Sports, Science and Technology (MEXT), Japan, Grant Number JPMXP09A19OS0052 at the Research Center for Ultra-High Voltage Electron Microscopy (Nanotechnology Open Facilities) in Osaka University and by the Collaborative Research Program of Institute for Protein Research, Osaka University, CEMCR-1701.

## Data availability

The filaments coordinates from this publication have been deposited in the Protein Data Bank (PDB, https://www.rcsb.org/) under accession codes are 8X1I (*Dh*-cParM1-ADP) and for *Cb*-cParM: 7X54, 7X56, 7X59 and 7X55 for ADP state, GDP state, the second class with GTP, and the short incubation time with GTP, respectively. The corresponding EM maps have been deposited in the Electron Microscopy Data Bank (EMDB, https://www.ebi.ac.uk/emdb/) are EMD-37996 (*Dh*-cParM1) and EMD-33007, 33009, 33012 and 33008 (*Cb*-cParM). The apo *Cb*-cParM X-ray structure is deposited in the PDB (7X3H). All other data are available from the corresponding author upon reasonable request.

## Competing Interest Statement

The authors declare no competing interest.

## Footnotes

† **Author Contributions**: These authors contributed equally to this work. AK, SA, RCR, and AN designed the study. AK, SA, and DP performed protein purification. SA and DP performed biochemical experiments and light microscopy. AK, SA, NM, KI, and AN performed electron microscopy. RCR solved the crystal structure. AK, SA and AN performed image analysis and model building. RCR performed crystallography. YK performed sequence analysis. AK, SA, RCR, and AN wrote the manuscript.

## Notes

### Competing Interest Statement

The authors have declared no competing interest.

### Summary of Updates

Movie S1 was added to depict the movement of the strand due to the phosphate release.

## References

Adams PD, Afonine P V, Bunkóczi G, Chen VB, Davis IW, Echols N, Headd JJ, Hung L-W, Kapral GJ, Grosse-Kunstleve RW. 2010. PHENIX: a comprehensive Python-based system for macromolecular structure solution. Acta Crystallogr D Biol Crystallogr 66:213–221.

Adams PD, Afonine P V, Bunkóczi G, Chen VB, Echols N, Headd JJ, Hung L-W, Jain S, Kapral GJ, Kunstleve RWG. 2011. The Phenix software for automated determination of macromolecular structures. Methods 55:94–106.

Adams V, Watts TD, Bulach DM, Lyras D, Rood JI. 2015. Plasmid partitioning systems of conjugative plasmids from Clostridium perfringens. Plasmid 80:90–96.

Bengelsdorf FR, Poehlein A, Esser C, Schiel-Bengelsdorf B, Daniel R, Dürre P. 2015. Complete genome sequence of the acetogenic bacterium Moorella thermoacetica DSM 2955T. Genome Announc 3:e01157–15.

Bharat TAM, Murshudov GN, Sachse C, Löwe J. 2015. Structures of actin-like ParM filaments show architecture of plasmid-segregating spindles. Nature 523:106–110.

Brzoska AJ, Jensen SO, Barton DA, Davies DS, Overall RL, Skurray RA, Firth N. 2016. Dynamic filament formation by a divergent bacterial actin-like ParM protein. PLoS One 11:e0156944.

Camacho C, Coulouris G, Avagyan V, Ma N, Papadopoulos J, Bealer K, Madden TL. 2009. BLAST+: architecture and applications. BMC Bioinformatics 10:1–9.

Campbell CS, Mullins RD. 2007. In vivo visualization of type II plasmid segregation: bacterial actin filaments pushing plasmids. J Cell Biol 179:1059–1066.

Carlier M-F, Laurent V, Santolini J, Melki R, Didry D, Xia G-X, Hong Y, Chua N-H, Pantaloni D. 1997. Actin depolymerizing factor (ADF/cofilin) enhances the rate of filament turnover: implication in actin-based motility. J Cell Biol 136:1307–1322.

Choi CL, Claridge SA, Garner EC, Alivisatos AP, Mullins RD. 2008. Protein-nanocrystal conjugates support a single filament polymerization model in R1 plasmid segregation. Journal of Biological Chemistry 283:28081–28086.

Croll TI. 2018. ISOLDE: a physically realistic environment for model building into low-resolution electron-density maps. Acta Crystallogr D Struct Biol 74:519–530.

Derman AI, Becker EC, Truong BD, Fujioka A, Tucey TM, Erb ML, Patterson PC, Pogliano J. 2009. Phylogenetic analysis identifies many uncharacterized actin-like proteins (Alps) in bacteria: regulated polymerization, dynamic instability and treadmilling in Alp7A. Mol Microbiol 73:534–552.

Ebersbach G, Gerdes K. 2005. Plasmid segregation mechanisms. Annu Rev Genet 39:453–479.

von der Ecken J, Heissler SM, Pathan-Chhatbar S, Manstein DJ, Raunser S. 2016. Cryo-EM structure of a human cytoplasmic actomyosin complex at near-atomic resolution. Nature 534:724–728.

Emsley P, Cowtan K. 2004. Coot: model-building tools for molecular graphics. Acta Crystallogr D Biol Crystallogr 60:2126–2132.

Emsley P, Lohkamp B, Scott WG, Cowtan K. 2010. Features and development of Coot. Acta Crystallogr D Biol Crystallogr 66:486–501.

Fogel MA, Waldor MK. 2005. Distinct segregation dynamics of the two Vibrio cholerae chromosomes. Mol Microbiol 55:125–136.

Fujiwara I, Vavylonis D, Pollard TD. 2007. Polymerization kinetics of ADP-and ADP-Pi-actin determined by fluorescence microscopy. Proceedings of the National Academy of Sciences 104:8827–8832.

Galkin VE, Orlova A, Rivera C, Mullins RD, Egelman EH. 2009. Structural polymorphism of the ParM filament and dynamic instability. Structure 17:1253–1264.

Garner EC, Campbell CS, Mullins RD. 2004. Dynamic instability in a DNA-segregating prokaryotic actin homolog. Science (1979) 306:1021–1025.

Garner EC, Campbell CS, Weibel DB, Mullins RD. 2007. Reconstitution of DNA segregation driven by assembly of a prokaryotic actin homolog. Science (1979) 315:1270–1274.

Gayathri P, Fujii T, Møller-Jensen J, Van Den Ent F, Namba K, Löwe J. 2012. A bipolar spindle of antiparallel ParM filaments drives bacterial plasmid segregation. Science (1979) 338:1334– 1337.

Gilchrist CLM, Chooi Y-H. 2021. Clinker & clustermap. js: Automatic generation of gene cluster comparison figures. Bioinformatics 37:2473–2475.

Goddard TD, Huang CC, Meng EC, Pettersen EF, Couch GS, Morris JH, Ferrin TE. 2018. UCSF ChimeraX: Meeting modern challenges in visualization and analysis. Protein Science 27:14–25.

He S, Scheres SHW. 2017. Helical reconstruction in RELION. J Struct Biol 198:163–176.

Jensen RB, Gerdes K. 1997. Partitioning of plasmid R1. The ParM protein exhibits ATPase activity and interacts with the centromere-like ParR-parC complex. J Mol Biol 269:505–513.

Jiang S, Narita A, Popp D, Ghoshdastider U, Lee LJ, Srinivasan R, Balasubramanian MK, Oda T, Koh F, Larsson M. 2016. Novel actin filaments from Bacillus thuringiensis form nanotubules for plasmid DNA segregation. Proceedings of the National Academy of Sciences 113:E1200–E1205.

Kabsch W, Holmes KC. 1995. The actin fold. The FASEB Journal 9:167–174.

Kabsch W, Mannherz HG, Suck D, Pai EF, Holmes KC. 1990. Atomic structure of the actin: DNase I complex. Nature 347:37–44.

Kadoya R, Baek JH, Sarker A, Chattoraj DK. 2011. Participation of chromosome segregation protein ParAI of Vibrio cholerae in chromosome replication. J Bacteriol 193:1504–1514.

Kimanius D, Dong L, Sharov G, Nakane T, Scheres SHW. 2021. New tools for automated cryo-EM single-particle analysis in RELION-4.0. Biochemical Journal 478:4169–4185.

Kodama T, Fukui K, Kometani K. 1986. The initial phosphate burst in ATP hydrolysis by myosin and subfragment-1 as studied by a modified malachite green method for determination of inorganic phosphate. The Journal of Biochemistry 99:1465–1472.

Koh F, Narita A, Lee LJ, Tanaka K, Tan YZ, Dandey VP, Popp D, Robinson RC. 2019. The structure of a 15-stranded actin-like filament from Clostridium botulinum. Nat Commun 10:1–10.

Lewis PJ, Errington J. 1997. Direct evidence for active segregation of oriC regions of the Bacillus subtilis chromosome and co-localization with the Spo0J partitioning protein. Mol Microbiol 25:945–954.

Li X, Mooney P, Zheng S, Booth CR, Braunfeld MB, Gubbens S, Agard DA, Cheng Y. 2013. Electron counting and beam-induced motion correction enable near-atomic-resolution single-particle cryo-EM. Nat Methods 10:584–590.

Livny J, Yamaichi Y, Waldor MK. 2007. Distribution of centromere-like parS sites in bacteria: insights from comparative genomics. J Bacteriol 189:8693–8703.

Mohl DA, Gober JW. 1997. Cell cycle–dependent polar localization of chromosome partitioning proteins in Caulobacter crescentus. Cell 88:675–684.

Møller-Jensen J, Borch J, Dam M, Jensen RB, Roepstorff P, Gerdes K. 2003. Bacterial mitosis: ParM of plasmid R1 moves plasmid DNA by an actin-like insertional polymerization mechanism. Mol Cell 12:1477–1487.

Narita A. 2011. Minimum requirements for the actin-like treadmilling motor system. Bioarchitecture 1:205–208.

Nonaka H, Keresztes G, Shinoda Y, Ikenaga Y, Abe M, Naito K, Inatomi K, Furukawa K, Inui M, Yukawa H. 2006. Complete genome sequence of the dehalorespiring bacterium Desulfitobacterium hafniense Y51 and comparison with Dehalococcoides ethenogenes 195. J Bacteriol 188:2262–2274.

Nordström K, Austin SJ. 1989. Mechanisms that contribute to the stable segregation of plasmids. Annu Rev Genet 23:37–69.

Oda T, Takeda S, Narita A, Maéda Y. 2019. Structural polymorphism of actin. J Mol Biol 431:3217–3228.

Otwinowski Z, Minor W. 1997. [20] Processing of X-ray diffraction data collected in oscillation modeMethods in Enzymology. Elsevier. pp. 307–326.

Pettersen EF, Goddard TD, Huang CC, Couch GS, Greenblatt DM, Meng EC, Ferrin TE. 2004. UCSF Chimera—a visualization system for exploratory research and analysis. J Comput Chem 25:1605–1612.

Polka JK, Kollman JM, Agard DA, Mullins RD. 2009. The structure and assembly dynamics of plasmid actin AlfA imply a novel mechanism of DNA segregation. J Bacteriol 191:6219– 6230.

Pollard TD, Borisy GG. 2003. Cellular motility driven by assembly and disassembly of actin filaments. Cell 112:453–465.

Popp D, Narita A, Lee LJ, Ghoshdastider U, Xue B, Srinivasan R, Balasubramanian MK, Tanaka T, Robinson RC. 2012. Novel actin-like filament structure from Clostridium tetani. Journal of Biological Chemistry 287:21121–21129.

Popp D, Xu W, Narita A, Brzoska AJ, Skurray RA, Firth N, Goshdastider U, Maéda Y, Robinson RC, Schumacher MA. 2010. Structure and filament dynamics of the pSK41 actin-like ParM protein: implications for plasmid DNA segregation. Journal of Biological Chemistry 285:10130–10140.

Ptacin JL, Lee SF, Garner EC, Toro E, Eckart M, Comolli LR, Moerner WE, Shapiro L. 2010. A spindle-like apparatus guides bacterial chromosome segregation. Nat Cell Biol 12:791–798.

Rohl CA, Strauss CEM, Misura KMS, Baker D. 2004. Protein structure prediction using RosettaMethods in Enzymology. Elsevier. pp. 66–93.

Rohou A, Grigorieff N. 2015. CTFFIND4: Fast and accurate defocus estimation from electron micrographs. J Struct Biol 192:216–221.

Salje J, Löwe J. 2008. Bacterial actin: architecture of the ParMRC plasmid DNA partitioning complex. EMBO J 27:2230–2238.

Scheres SHW. 2012. RELION: implementation of a Bayesian approach to cryo-EM structure determination. J Struct Biol 180:519–530.

Senior AW, Evans R, Jumper J, Kirkpatrick J, Sifre L, Green T, Qin C, Žídek A, Nelson AWR, Bridgland A. 2020. Improved protein structure prediction using potentials from deep learning. Nature 577:706–710.

Szewczak-Harris A, Löwe J. 2018. Cryo-EM reconstruction of AlfA from Bacillus subtilis reveals the structure of a simplified actin-like filament at 3.4-Å resolution. Proceedings of the National Academy of Sciences 115:3458–3463.

Tanaka K, Takeda S, Mitsuoka K, Oda T, Kimura-Sakiyama C, Maéda Y, Narita A. 2018. Structural basis for cofilin binding and actin filament disassembly. Nat Commun 9:1860.

van den Ent F, Amos LA, Löwe J. 2001. Prokaryotic origin of the actin cytoskeleton. Nature 413:39–44.

van den Ent F, Møller-Jensen J, Amos LA, Gerdes K, Löwe J. 2002. F-actin-like filaments formed by plasmid segregation protein ParM. EMBO J 21:6935–6943.

Wang H, Robinson RC, Burtnick LD. 2010. The structure of native G-actin. Cytoskeleton 67:456– 465.

Wegner A. 1976. Head to tail polymerization of actin. J Mol Biol 108:139–150.

Wickstead B, Gull K. 2011. The evolution of the cytoskeleton. Journal of Cell Biology 194:513– 525.

Yasunaga T, Wakabayashi T. 1996. Extensible and object-oriented system Eos supplies a new environment for image analysis of electron micrographs of macromolecules. J Struct Biol 116:155–160.

Zhang K. 2016. Gctf: Real-time CTF determination and correction. J Struct Biol 193:1–12.

Zhao B, Mesbah NM, Dalin E, Goodwin L, Nolan M, Pitluck S, Chertkov O, Brettin TS, Han J, Larimer FW. 2011. Complete genome sequence of the anaerobic, halophilic alkalithermophile Natranaerobius thermophilus JW/NM-WN-LF.

Zivanov J, Nakane T, Forsberg BO, Kimanius D, Hagen WJH, Lindahl E, Scheres SHW. 2018. New tools for automated high-resolution cryo-EM structure determination in RELION-3. Elife 7:e42166.

