## Supplementary data for "Bacterial genome encoded ParMs"

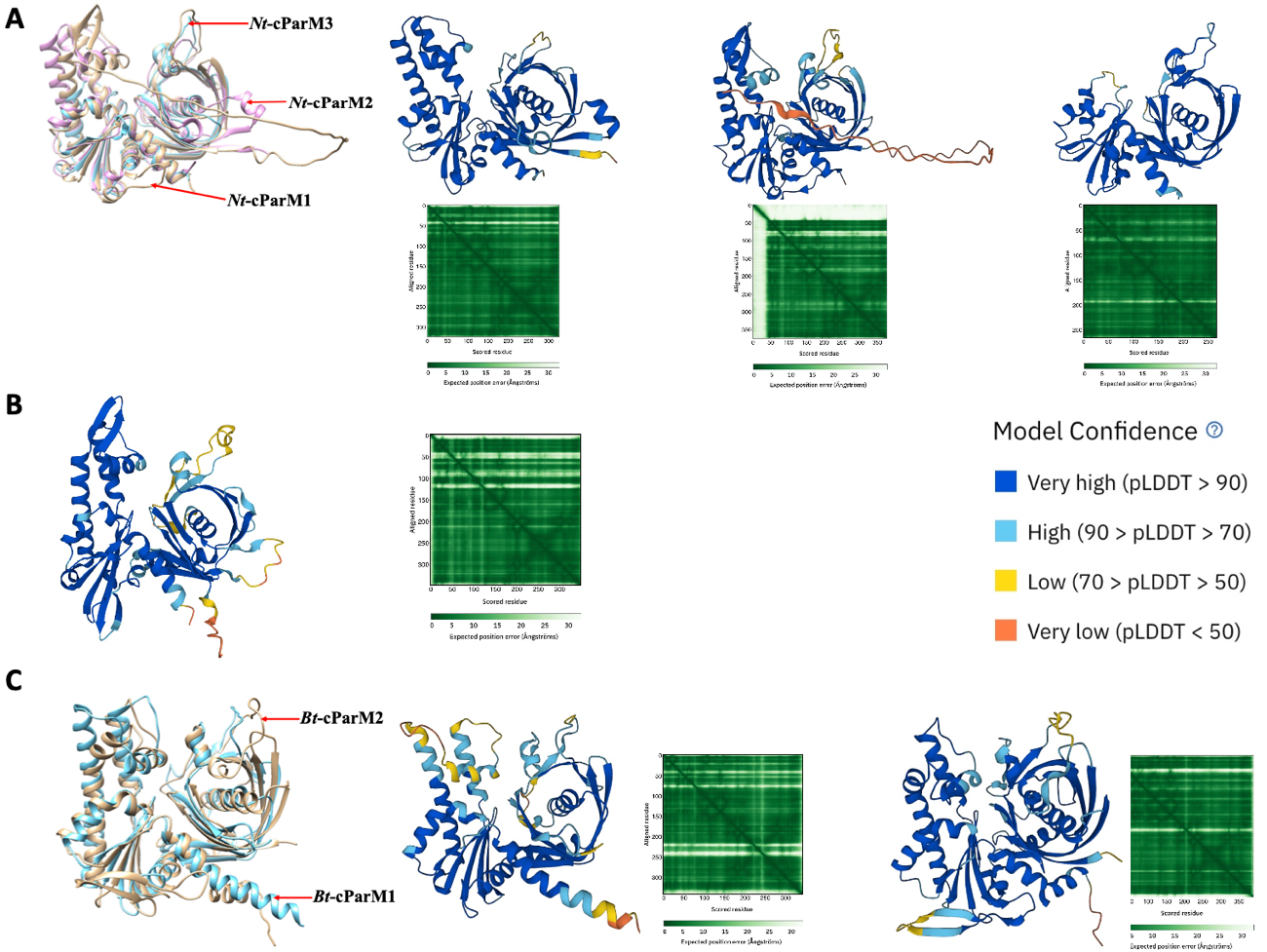

**Figure S1: AF2 models of selected cParMs. (A)** *Nt*-cParMs – *Nt*-cParM1 (WP_148206872.1) (pink), *Nt*-cParM2 (WP_012448769.1) (brown) and *Nt*-cParM3 (WP_012446843.1) (blue) encoded on the *Natranaerobius thermophilus* JW/NM-WN-LF strain chromosome **(B**) A single copy of *Mt*-cParM (WP_011391888.1) encoded on the chromosome from *Moorella thermoacetica* strain 39073-HH **(C)** *Bt*-cParMs - *Bt*-cParM1 (WP_001968526.1) (blue) and *Bt*-cParM2 (WP_000025611.1) (brown) encoded on the *Bacillus tropicus* strain FDAARGOS_920 chromosome. Each individual AF2 model is shown with different colours based on the pLDDT model confidence score. For pLTTD plots, AlphaFold2 produces a per-residue model confidence score (pLDDT) between 0 and 100. Some regions below 50 pLDDT may be unstructured in isolation. Below or adjacent the models are the respective Predicted Aligned Error (PAE) plots of the various regions of the models. Each 2D plot square shade of green indicates the expected distance error in Ångströms (Å) for a pair of residues. A dark green tile designates a good prediction or low error whereas a light green tile indicates poor prediction or high error.

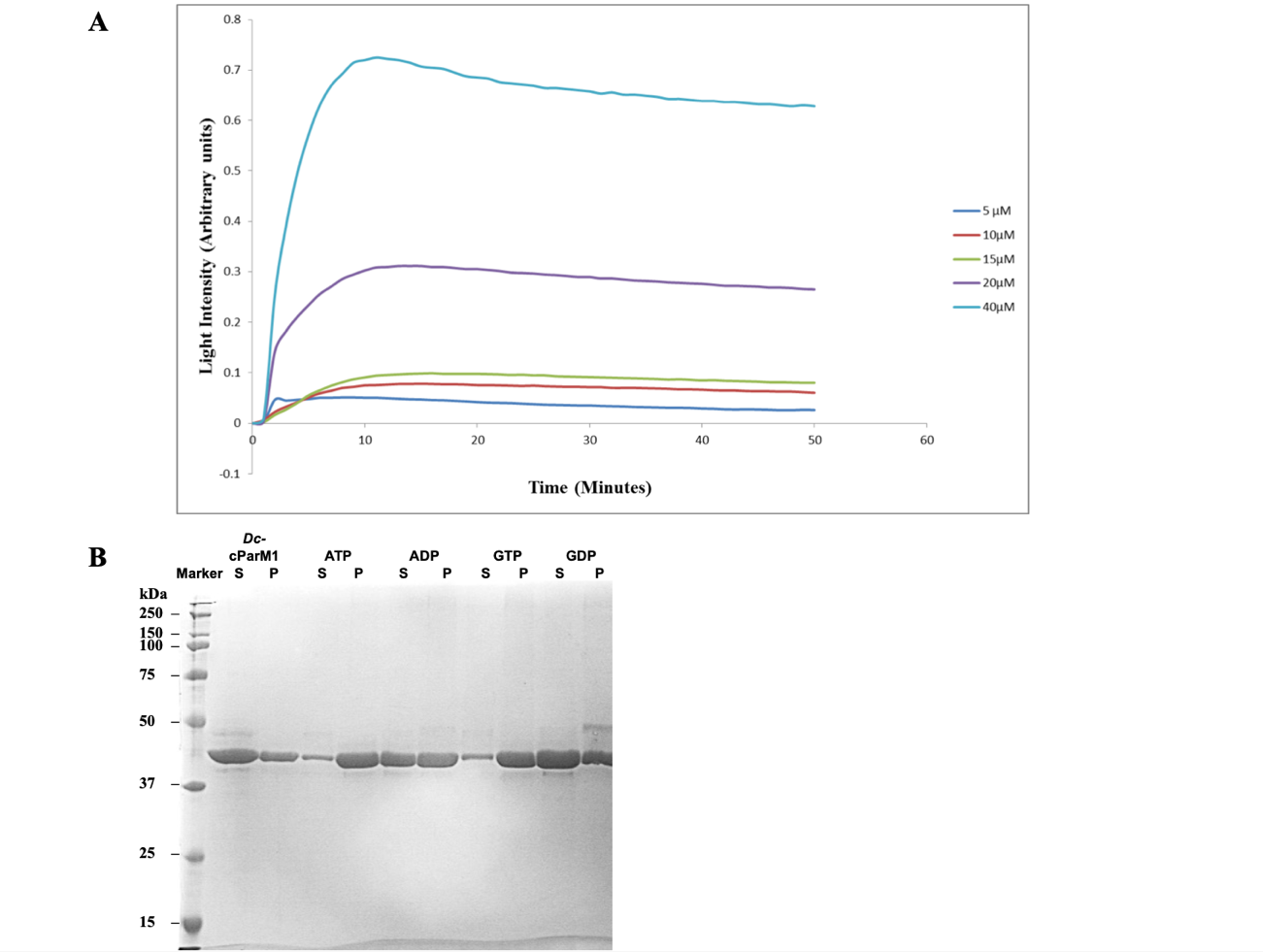

**Figure S2: *Dh*-cParM1 polymerization with different nucleotides. (A)** Different *Dh*-cParM1 concentrations polymerized with 5 mM ATP monitored by light scattering. All readings were taken at a wavelength of 600 nm and in a potassium chloride buffer (300 mM potassium chloride, 40 mM HEPES, pH 7.5, 1 mM MgCl_2_) at 25 °C **(B)** An SDS-PAGE gel of the sedimentation assay indicating the polymerization of 20 µM *Dh*-cParM1 with/without 5 mM of different nucleotides. Key: Marker, the standard protein marker; *Dh*-cParM1, protein without nucleotide; other lanes indicate the nucleotide used; S, soluble fraction; P, pellet fraction.

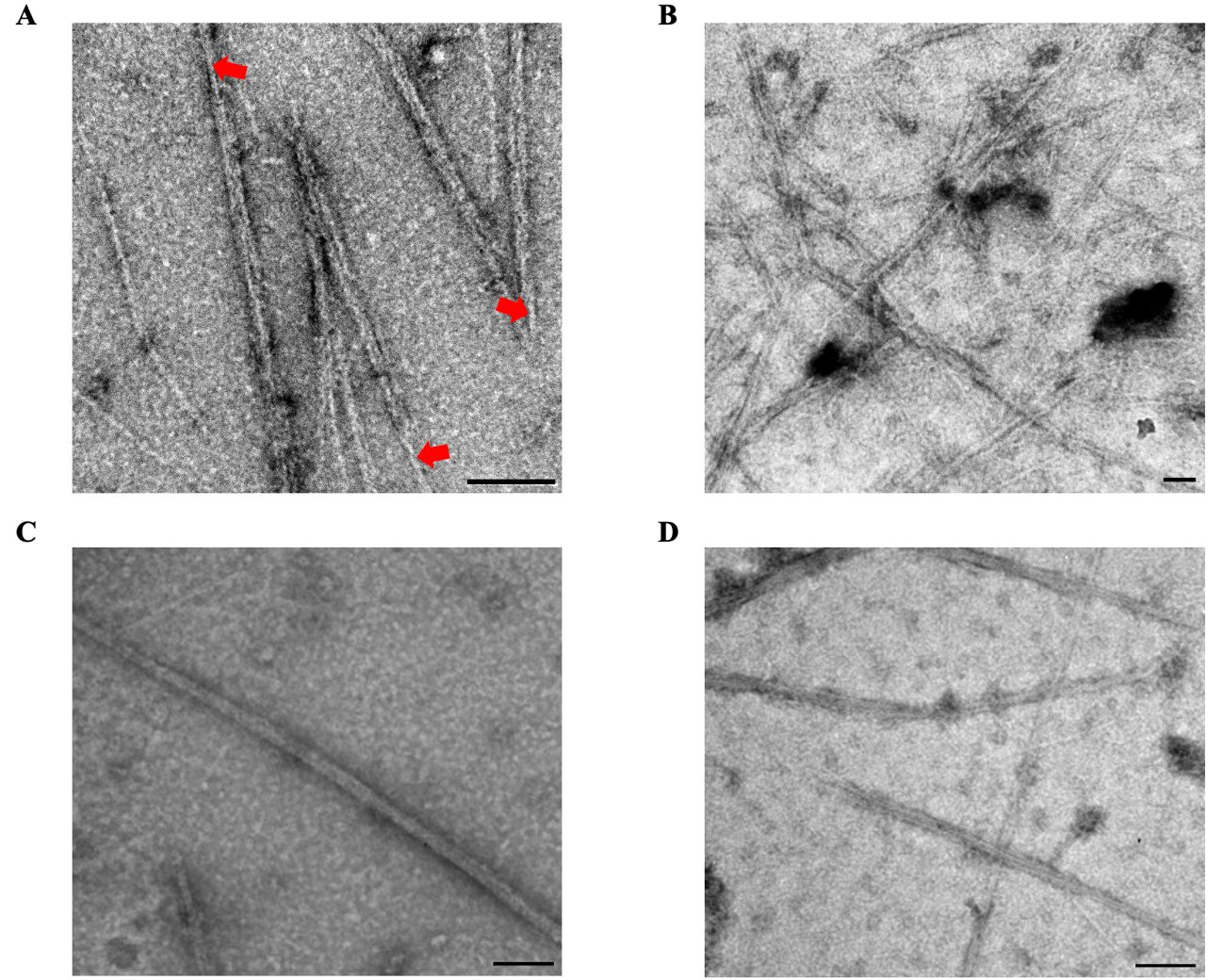

**Figure S3: *Dh*-cParM1 confirmation of filament formation in negatively stained samples.** 20 µM *Dh*-cParM1 polymerized with 5 mM of different nucleotides. Micrographs displaying the two types of *Dh*-cParM1 filament morphologies formed by (**A**) ATP (**B**) GTP (**C** ) AMPPNP (**D**) ATP-γ-S. The widths of coupled filaments narrow at the ends of the filament as indicated by the red arrow heads in **(A)**. All micrographs were prepared in potassium chloride buffer (300 mM potassium chloride, 40 mM HEPES, pH 7.5, 1 mM MgCl_2_) incubated at 25 °C. Scalebar = 100 nm.

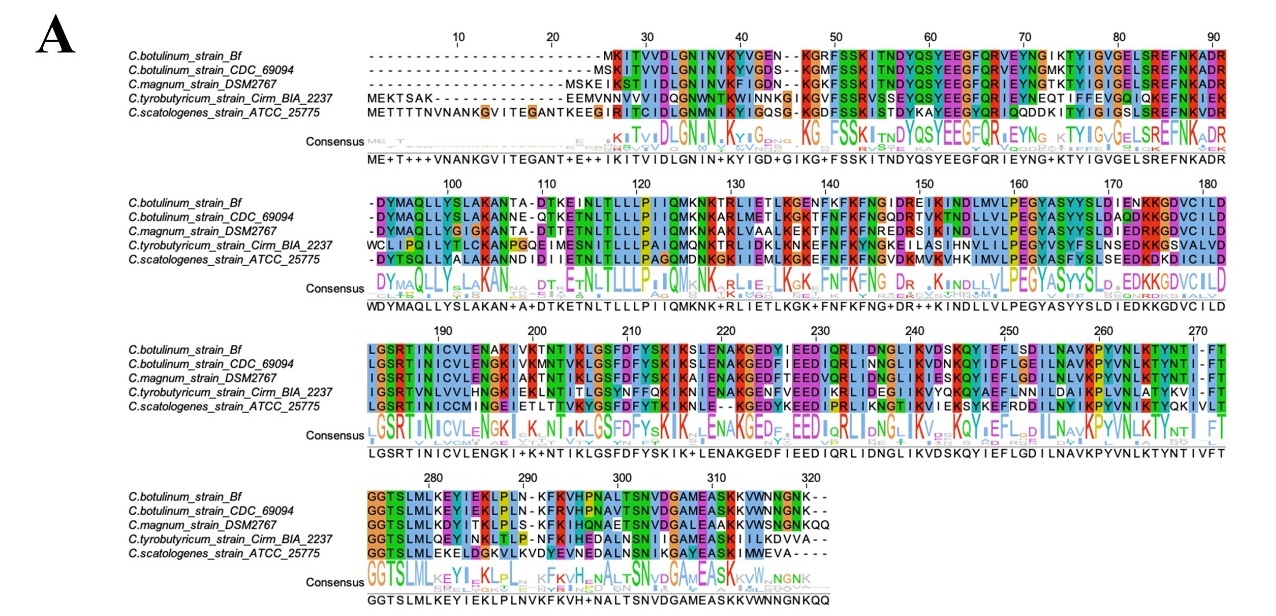

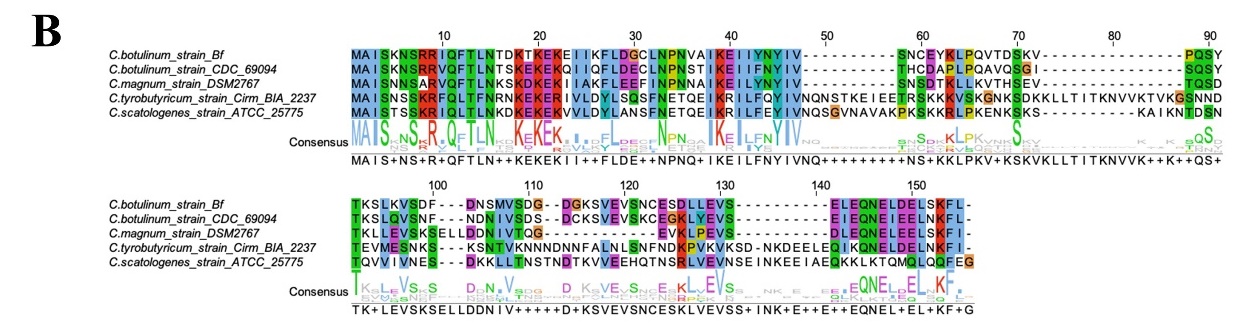

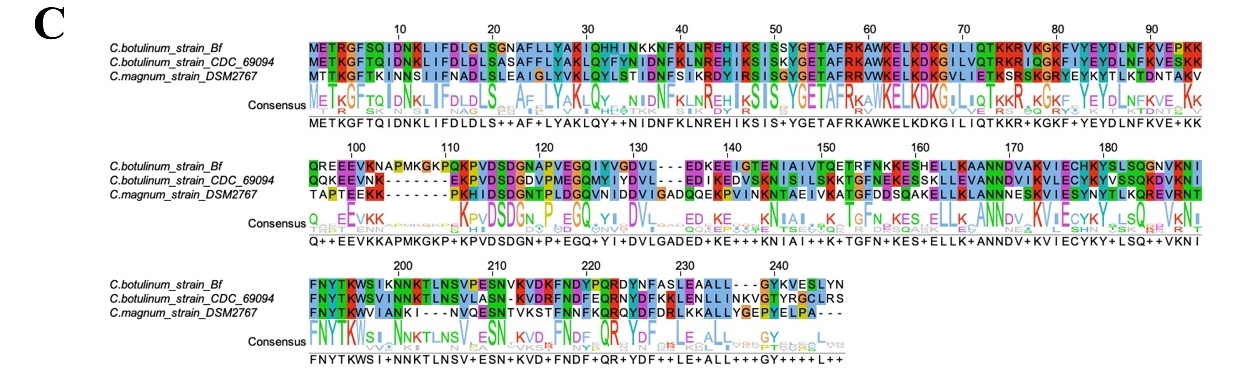

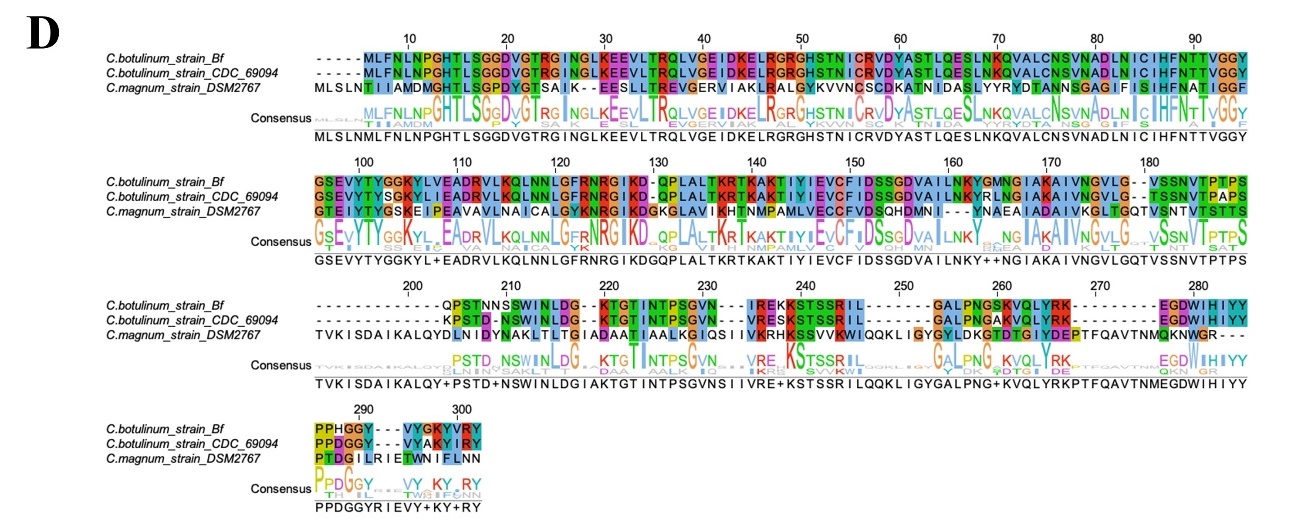

**Figure S4: Multiple sequence alignments of homologous sequences of cParMRC gene clusters in *Clostridium sp*.** Multiple sequence alignment of cParM **(A)**, cParR (B), putative replication initiator **(C),** and sporulation-specific N-acetylmuramoyl-L-alanine amidase **(D)** (one of the genes) were performed with MUSCLE (Edgar, 2004). Homologous residues are colored according to the ClustalX scheme.

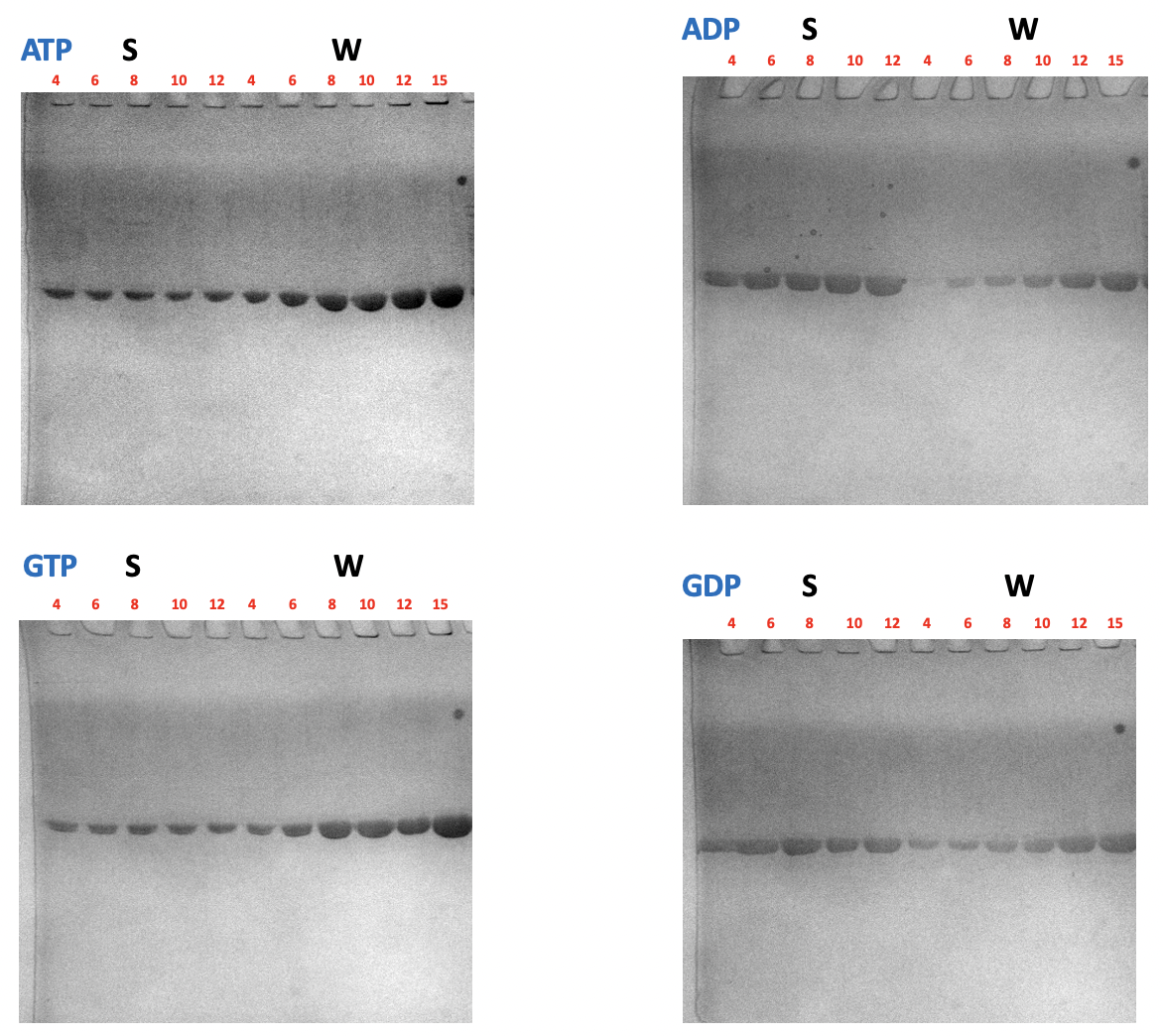

**Figure S5. Sedimentation assay for estimating *Cb*-cParM** **critical concentration induced by different nucleotides.** S represents supernatant and W represents whole before sedimentation. The number above each lane represents the concentration of *Cb*-cParM (µM). *Cb*-cParM concentrations from W lanes were used to plot a standard curve to estimate the *Cb*-cParM concentrations of S lanes.

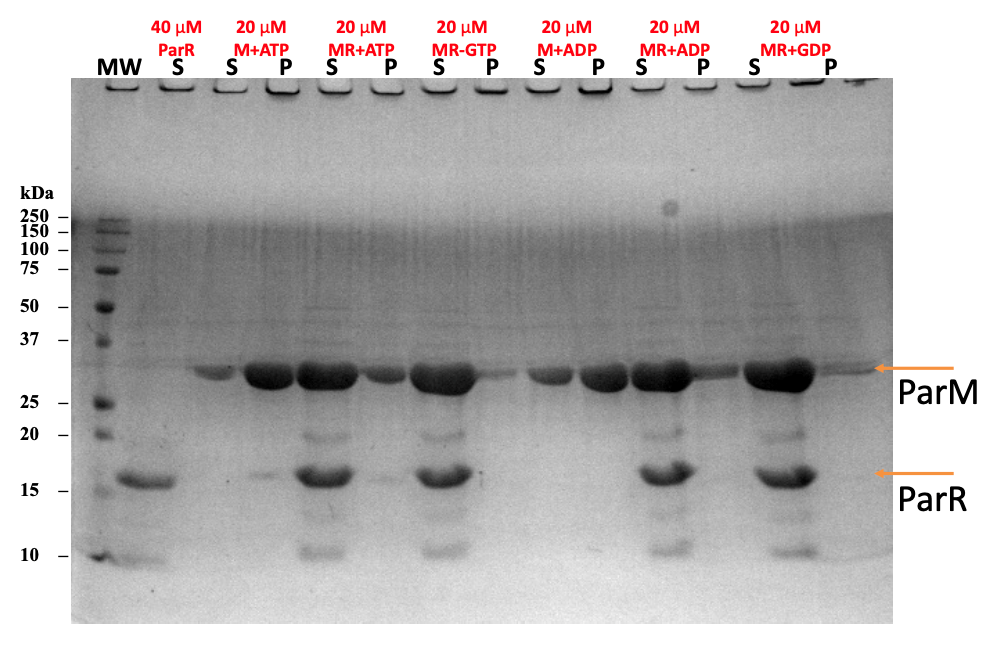

**Figure S6: Sedimentation assay for *Cb*-cParM with and without *Cb*-cParR.** M and MR represent *Cb*-cParM without or with *Cb*-cParR, respectively. S and P represent supernatant and pellet, respectively. The clear decrease in *Cb*-cParM concentration in the pellet fraction with *Cb*-cParR indicates the depolymerizing function of *Cb*-cParR.

**References**

Edgar RC (2004) MUSCLE: multiple sequence alignment with high accuracy and high throughput. *Nucleic Acids Res* 32: 1792-1797

**Movie S1: Structural shift in *Cb*-cParM filaments.** Models and maps for the class1 (corresponding to GDP state) and the class2 with GTP were aligned by the ID rigid bodies of the center subunit.

**Table S1. Percentage sequence identities of chromosome encoded cParMs**

|  | cParM1  *Natranaerobius*  *thermophilus* | cParM-2  *Natranaerobius thermophilus* | cParM3  *Natranaerobius thermophilus* | *Caloramator* sp | *Clostridium sp* | *Bacillus sp* |
| --- | --- | --- | --- | --- | --- | --- |
| cParM1  *Natranaerobius thermophilus* | 100.0 | 24.9 | <15.0 | 44.6 | 37.8 | 27.5 |
| cParM2  *Natranaerobius thermophilus* | 24.9 | 100.0 | 31.0 | 23.0 | 21.0 | <15.0 |
| cParM3  *Natranaerobius thermophilus* | <15.0 | 31.0 | 100.0 | 23.2 | 26.7 | 27.5 |
| *Caloramator sp* | 44.6 | 23.0 | 23.2 | 100.0 | 46.2 | 30.3 |
| *Clostridium sp* | 37.8 | 21.0 | 26.7 | 46.2 | 100.0 | 27.1 |
| *Bacillus sp* | 27.5 | <15.0 | 27.5 | 30.3 | 27.1 | 100.0 |

**Table S2. Percentage sequence identities of *Moorella* chromosome encoded cParMs**

|  | *Moorella thermoacetica* | *Moorella* sp. | *Moorella glycerini* | *Thermoanaeo*  *bacteraceae* | *Moorella humiferrea* | *Moorella mulderi* |
| --- | --- | --- | --- | --- | --- | --- |
| *Moorella thermoacetica* | 100.0 | 95.7 | 94.8 | 92.5 | 94.2 | 94.2 |
| *Moorella* sp. | 95.7 | 100.0 | 93.7 | 92.0 | 96.8 | 93.1 |
| *Moorella glycerini* | 94.8 | 93.7 | 100.0 | 92.8 | 92.8 | 93.7 |
| *Thermoanaeo*  *bacteraceae* | 92.5 | 92.0 | 92.8 | 100.0 | 90.8 | 91.1 |
| *Moorella humiferrea* | 94.2 | 96.8 | 92.8 | 90.8 | 100.0 | 93.1 |
| *Moorella mulderi* | 94.2 | 93.1 | 93.7 | 91.1 | 93.1 | 100.0 |

**Table S3. Percentage sequence identities of *Bacillus* chromosome encoded cParMs**

|  | cParM1  *Bacillus tropicus* | cParM2  *Bacillus tropicus* | *Bacillus cereus* | *Bacillus anthracis* | *Bacillus paranthracis* | *Bacillus pacificus* |
| --- | --- | --- | --- | --- | --- | --- |
| cParM1  *Bacillus tropicus* | 100.0 | <15.0 | 99.1 | 98.5 | 97.1 | 95.9 |
| cParM2  *Bacillus tropicus* | <15.0 | 100.0 | <15.0 | <15.0 | <15.0 | <15.0 |
| *Bacillus cereus* | 99.1 | <15.0 | 100.0 | 98.8 | 96.5 | 96.5 |
| *Bacillus anthracis* | 98.5 | <15.0 | 98.8 | 100.0 | 96.5 | 96.5 |
| *Bacillus paranthracis* | 97.1 | <15.0 | 96.5 | 96.8 | 100.0 | 99.4 |
| *Bacillus pacificus* | 97.1 | <15.0 | 96.5 | 96.5 | 99.4 | 100.0 |

**Table S4: Statistics of *Dh*-cParM1 protomer, cryoEM and model refinement data**

|  | *Dh*-cParM1 (EMD-37996) (PDB 8X1I) |
| --- | --- |
| **Data collection and processing** |  |
| Voltage (kV) | 300 kV |
| Electron exposure (e–/Å^2^) | 40.0 |
| Defocus range (μm) | ∼ −1.5 −2.3 |
| Pixel size (Å) | 1.1 |
| Phase plate | No |
| Symmetry imposed | Helical |
| Final particle images (no.) | 71060 |
| Map resolution (Å)  FSC threshold | 4.0  0.143 |
| Map resolution range (Å) | ∞ ~ 4.0 |
| **Refinement** |  |
| Initial model used (PDB code) | 8X1I |
| Map sharpening *B* factor (Å^2^) | -241 |
| Model composition  Non-hydrogen atoms  Protein residues  Ligands | 2882  369  2 |
| R.m.s. deviations  Bond lengths (Å)  Bond angles (°) | 0.012  1.973 |
| Validation  MolProbity score  Clashscore  Poor rotamers (%) | 1.21  0.0  3 |
| Ramachandran plot  Favored (%)  Allowed (%)  Disallowed (%) | 95  5  0 |

**Table S5. Cryo-EM data collection, refinement, and validation statistics**

|  | # ADP  (EMDB-33007)  (PDB 7X54) | #2 GDP  (EMDB-33009)  (PDB 7X56) | #3 GTP class 2  (EMDB-33012)  (PDB 7X59) | #4 GTP short incubation  (EMDB-33008)  (PDB 7X55) |
| --- | --- | --- | --- | --- |
| **Data collection and processing** |  |  |  |  |
| Voltage (kV) | 300 kV | 300 kV | 300 kV | 300 kV |
| Electron exposure (e–/Å^2^) | 45 | 45 | 45 | 45 |
| Defocus range (μm) | -1.5 ~ -3.5 | -1.0 ~ -3.0 | -1.0 ~ -3.0 | -0.5 ~ -1.5 |
| Pixel size (Å) | 0.87 | 0.87 | 0.87 | 0.87 |
| Phase plate | No | No | No | Yes |
| Symmetry imposed | Helical | Helical | Helical | Helical |
| Final particle images (no.) | 36762 | 40599 | 70754 | 153326 |
| Map resolution (Å)  FSC threshold | 3.9  0.143 | 3.5  0.143 | 6.5  0.143 | 8.6  0.143 |
| Map resolution range (Å) | ∞ ~ 3.9 | ∞ ~ 3.5 | ∞ ~ 6.5 | ∞ ~ 8.6 |
| **Refinement** |  |  |  |  |
| Initial model used (PDB code) | 7X56 | 6IZV | 7X56 | 7X56 |
| Map sharpening *B* factor (Å^2^) | -105 | -117 | -527 | -1281 |
| Model composition  Non-hydrogen atoms  Protein residues  Ligands | 11575  1425  10 | 11580  1425  10 | 11595  1425  5 | 11600  1425  5 |
| R.m.s. deviations  Bond lengths (Å)  Bond angles (°) | 0.004  0.979 | 0.005  0.965 | 0.004  1.059 | 0.007  1.137 |
| Validation  MolProbity score  Clashscore  Poor rotamers (%) | 1.98  8.79  0.39 | 1.96  7.71  0.00 | 2.19  11.95  0 | 2.44  21.32  0 |
| Ramachandran plot  Favored (%)  Allowed (%)  Disallowed (%) | 91  9  0 | 90  10  0 | 88  12  0 | 89  11  0 |

**Table S6. X-ray crystallography data collection, refinement and validation statistics**

|  | *Cb*-cParM  (PDB code 7X3H) |
| --- | --- |
| **Protein**  Accession No.  Mutations | EDT87363.1  R204D, K230D, N234D |
| **Crystal** | P2_1_ |
| *a*, *b*, *c* (Å) | 55.6, 51.1, 64.9 |
| *α, β, γ* (°) | 90.0, 115.3, 90.0 |
| **Data collection** |  |
| Wavelength (Å) | 1.0 |
| Resolution (Å)^a^ | 50.0-1.7 (1.73-1.70) |
| *R*_merge_ | 3.0 (47.3) |
| *R*_meas_ | 3.5 (58.8) |
| *R*_pim_ | 1.8 (34.4) |
| *I/*σ(*I*) | 37.7 (1.8) |
| *CC*_1/2_ | (0.736) |
| Completeness (%) | 99.3 (94.1) |
| Redundancy | 3.6 (2.5) |
| **Refinement** |  |
| Resolution (Å) | 32.0-1.7 (1.76-1.70) |
| No. reflections | 35734 (2896) |
| *R*_work_ / *R*_free_ | 19.2/22.4 (27.1/29.9) |
| No. atoms  Protein | 2249 |
| Water | 314 |
| *B* factors |  |
| Protein | 25.4 |
| Water | 34.0 |
| r.m.s deviations |  |
| Bond lengths (Å) | 0.007 |
| Bond angles (°) | 1.09 |
| Ramachandran Plot |  |
| Favoured (%) | 98.5 |
| Outliers (%) | 0 |
